# Reverse-Engineering the Cortical Architecture for Controlled Semantic Cognition

**DOI:** 10.1101/860528

**Authors:** Rebecca L. Jackson, Timothy T. Rogers, Matthew A. Lambon Ralph

**Author notes:** Correspondence to: Dr. Rebecca Jackson, MRC Cognition & Brain Sciences Unit, University of Cambridge, 15 Chaucer Road, Cambridge, CB2 7EF.

## Abstract

We present a ‘reverse engineering’ approach to deconstruct cognition into neurocomputational mechanisms and their underlying cortical architecture, using controlled semantic cognition as a test case. By systematically varying the structure of a computational model and assessing the functional consequences, we identified architectural properties necessary for generating the core functions of the semantic system. Semantic cognition presents a challenging test case as the brain must achieve two seemingly contradictory functions: abstracting context-invariant conceptual representations across time and modalities, whilst producing specific context-sensitive behaviours appropriate for the immediate task. These functions were best achieved in models possessing a single, deep multimodal hub with sparse connections from modality-specific inputs, and control systems acting on peripheral rather than deep network layers. These architectural features correspond well with those suggested by neural data, strongly supporting the efficacy of the reverse engineering approach, and further generating novel hypotheses about the neuroanatomy of controlled semantic cognition.

---

At heart, cognitive neuroscience is an effort to understand how mental representations and processes arise from, and relate to, underlying neural mechanisms. Toward this goal, researchers typically begin by seeking relationships between neural data (neurophysiological activity and/or neuropathology) and patterns of behaviour in various tasks. We introduce an alternative ‘reverse engineering’ approach that first considers the functions a given cognitive system must support and then evaluates what neuro-computational machinery best achieves those functions. By systematically varying structure within a computer simulation and assessing the functional consequences, it is possible to establish the architectural elements critical to the targeted functions. These important elements provide a yardstick for assessing the actual architecture of the neural system under investigation, potentially explaining why the system is organised in a particular way.

We apply this approach to understand the cortical network underlying *semantic cognition*, the subdomain of cognition that supports controlled access to and manipulation of conceptual knowledge or meaning^1,2^. Semantic cognition provides a challenging test case because the system must concurrently achieve two functions that appear diametrically opposed. First, it must abstract over episodes and across time to acquire *context-independent representations* that express conceptual similarity structure and thereby promote knowledge generalisation across items and contexts^1,3,4^. This *conceptual abstraction* supports the ability to discern conceptual similarity relationships amongst items denoted by images, words, or other attributes, despite sometimes dramatic variability in their surface properties^4-6^. For instance, if one learns that wolves are dangerous after being attacked in the woods, this knowledge should generalise to a different wolf on a farm or in the house. The same conceptual representation should promote the inference ‘dangerous’ for all wolves across many situations. Second, the system must flexibly adapt semantic representations to suit immediate task demands^1,5,7,8^. For instance, differentiating wolves and dogs when generating an inference about safety, but treating them similarly for an inference about physical appearance. Thus, the subset of features governing the similarity space and consequent generalisation in the moment must also be *context-sensitive.*

Whilst the literature currently advances several hypotheses about the cortical architecture of the semantic system, no prior work has formally assessed their ability to simultaneously achieve conceptual abstraction and context-sensitivity. Interestingly, the extant hypotheses are variants of a central idea dating back at least to Wernicke: that semantic knowledge arises from interactions amongst various *surface representations* (sensory, motor, linguistic, affective, etc.) distributed throughout cortex^9^. They differ principally in their proposals about the pathways through which these various surface representations interact. Consequently, they can be contrasted using computer simulations with a family of neural network models, all learning to compute the same interactive mappings amongst various surface representations, but differing in their architecture. Illuminating the architectural elements that best support both conceptual abstraction and context sensitivity, generates a cognitively and computationally motivated hypothesis about cortical network structure that can be weighed against empirical evidence. Critical evidence includes extant data about the anatomy of the cortical semantic system as well as patterns of disordered behaviour in two contrasting semantic syndromes: semantic dementia (SD) - the incremental degradation of conceptual representations associated with gradual neuronal loss within the anterior temporal lobes^10-12^; and semantic aphasia (SA) - an impairment of semantic control following cerebrovascular accident to prefrontal or temporoparietal regions^2,13^. The rest of this paper reports such an analysis, arriving at a new model of controlled semantic cognition that explains the known anatomy of the cortical semantic system, and providing a template for reverse engineering in cognitive neuroscience, more generally.

## Core functions for semantic cognition

Research in semantic cognition has largely focused on conceptual abstraction: how do we acquire representations that capture conceptual structure from sensory, motor, and linguistic inputs that do not transparently reflect such structure? This ability is thought to arise from sensitivity to patterns of covariation in experience^14^. While birds vary wildly in appearance, they possess properties that covary together: feathers, beaks, wings, flying ability, the name “bird,” etc. Many otherwise competing theories agree that the human semantic system detects and exploits such structure, representing items as conceptually similar when they share coherently-covarying properties, even if they differ in many other respects^6,15,16^. By this view, concepts reflect clusters in the high-order covariance structure of experience.

This idea becomes challenging, however, when one considers how the various properties purported to co-occur in experience are distributed across learning episodes. While birds typically possess the abilities to fly and lay eggs, those behaviours do not directly co-occur: a bird laying an egg is not flying, and vice versa^6^. In general, each experience provides only partial exposure to an item’s properties. Moreover, such exposure can be highly context- and modality-specific; for instance, the fact that birds have hollow bones may only be encountered via verbal statements in science class. Thus, conceptual abstraction relies upon extracting the relevant covariances across many different episodes over time, each providing only limited, context-bound access to a subset of properties: the system must track sameness in kind across different contexts and experiences and detect that the flying item observed in one episode is similar to an item possessing feathers or labelled “bird” in another.

This requirement to form representations abstracted across items, modalities and contexts seems at odds with the second core function of semantic cognition, context-sensitivity. Context-appropriate behaviour requires flexible construction of a task-relevant similarity structure, as different subsets of features and aspects of meaning are crucial in different contexts, whilst other often more dominant meanings must be inhibited^1,6^. For instance, the system must represent a piano and a computer keyboard as similar when generating action plans, but as dissimilar when generating inferences about their weight. This flexibility underlies the construction of novel categories, such as objects that fit in a pocket^17^, and children’s qualitatively different generalisation patterns depending upon the nature of the generalised property^18^. Often the context-appropriate behaviour requires access to a particular subset of features in a desired sensory modality. For instance, the features relevant for moving a piano conflict with those relevant to playing it; accessing both simultaneously may produce an action inappropriate in strength or nature^7^. Equally, naming a piano requires accessing a highly specific verbal output from a different subset of information than miming its action, or drawing a piano. Thus, conceptual abstraction and context-sensitivity may appear diametrically opposed by virtue of their treatment of context, yet the semantic system must achieve both concurrently. How these antagonistic, yet highly interactive, processes are performed in tandem is a central unresolved issue in semantics research.

These considerations highlight three core functions of the human semantic system that informed the design of our simulations:

1. It must acquire representations that capture overall conceptual similarity structure and not merely the perceptual, motor, and linguistic structure apparent within various modalities.
2. It must acquire context-independent conceptual representations from learning episodes that provide only partial context-specific information about an item’s properties.
3. It must adapt to context so as to generate only context-appropriate behaviours.

## Proposals about the neural architecture of semantic representation

To our knowledge the Rumelhart model^19^ exhaustively analysed by Rogers and McClelland^6^ provides the only computational mechanism explaining how all three functions might be simultaneously realised; however, its feedforward structure makes it a poor candidate for understanding the cortical architecture. Hypotheses about the architecture of the semantic network abound, but few have been specified with sufficient mechanistic precision to compare and contrast their implications for understanding conceptual abstraction and context-sensitivity. The reverse-engineering approach we develop identifies a series of architectural *‘building blocks’* that distinguish various theories, then parametrically varies these to delineate a large space of possible architectures that encompass some existing proposals, as well as novel hypotheses not previously considered. Formal comparison of the effects of each building block provides critical insight into hypotheses that have been articulated only verbally in the literature, and allows us to determine which in the space of possible models best supports both conceptual abstraction and context sensitivity.

To identify the building blocks, we first considered how contemporary hypotheses about the cortical architecture of semantics vary. One view derives from Wernicke’s^9^ proposal that semantic processing reflects direct, interactive communication amongst various associated surface representations—perceptual, motor, linguistic, affective, and so on. This perspective foreshadows modern ‘embodied cognition’ views^20,21^ and has been explored computationally^22-25^. Other work emphasises the importance of multimodal ‘hub’ regions for mediating interactions between sensorimotor modalities. For instance, ‘convergence zones’ may connect different modalities within a network that adopts multiple hubs, such as a pathway connecting visual and linguistic representations, another connecting visual and haptic representations, a third connecting linguistic and haptic representations, and so on^26,27^. Such hubs could be the sole vehicle for cross-modal communication, or alternatively they could themselves connect via a broader multimodal region, producing a hierarchical convergence of information across modalities. Whilst neither proposal has been investigated computationally, both have received support from functional brain imaging and neuropsychological evidence^26,28,29^. Alternatively, a single region may be postulated to connect to, and mediate between, all sensory modalities, resulting in a single multimodal hub region. Finally, the various representational modalities might all communicate via a single multimodal hub, an idea supported by convergent computational^30,31^, neuropsychological^1,10,11^, neuroimaging^32,33^, neurophysiological^34,35^ and neurostimulation^36,37^ evidence.

With this landscape in mind, we considered two architectural factors that influence how modality-specific “spokes” connect to multimodal “hubs”. First, such communication could be direct or involve one or more intermediating regions. Deeper models (possessing more layers between input and output) can acquire more complex representations and behaviours as attested by the recent explosion of research in deep neural networks^38,39^, and several recent proposals suggest that conceptual representations arise within the deepest layers of visual and language networks^40-42^. Thus, depth seems an important factor to consider. Second, network layers may connect only to immediately adjacent areas, or may additionally send direct ‘shortcut’ connections to anatomically distal regions. Such shortcut connections have not previously been considered in semantic models, but provide an analogue of long-range connectivity supported by white matter pathways. Accordingly, such connectivity would need to be sparse due to metabolic and packing constraints within the brain^43,44^.

From these considerations we discern five building blocks that may influence the behaviour of the cortical semantic network: a) presence of multiple hubs connecting subsets of modality-specific spokes, b) presence of a single multimodal hub, c) network depth, d) presence of shortcut connections, and e) hierarchical convergence across modalities. These building blocks combine in different ways to produce the space of candidate architectures shown in Figure 1. Importantly, all architectures employ identical inputs/outputs, and possess equal numbers of hidden units and connections. They differ only in the pattern of connectivity amongst units, and hence in the organisation of units into layers and pathways. The central questions are how each building block impacts conceptual abstraction, first without and then with simultaneous context sensitivity, and whether successful architectures provide insight into existing anatomical and neuropsychological data.

**Figure 1.**
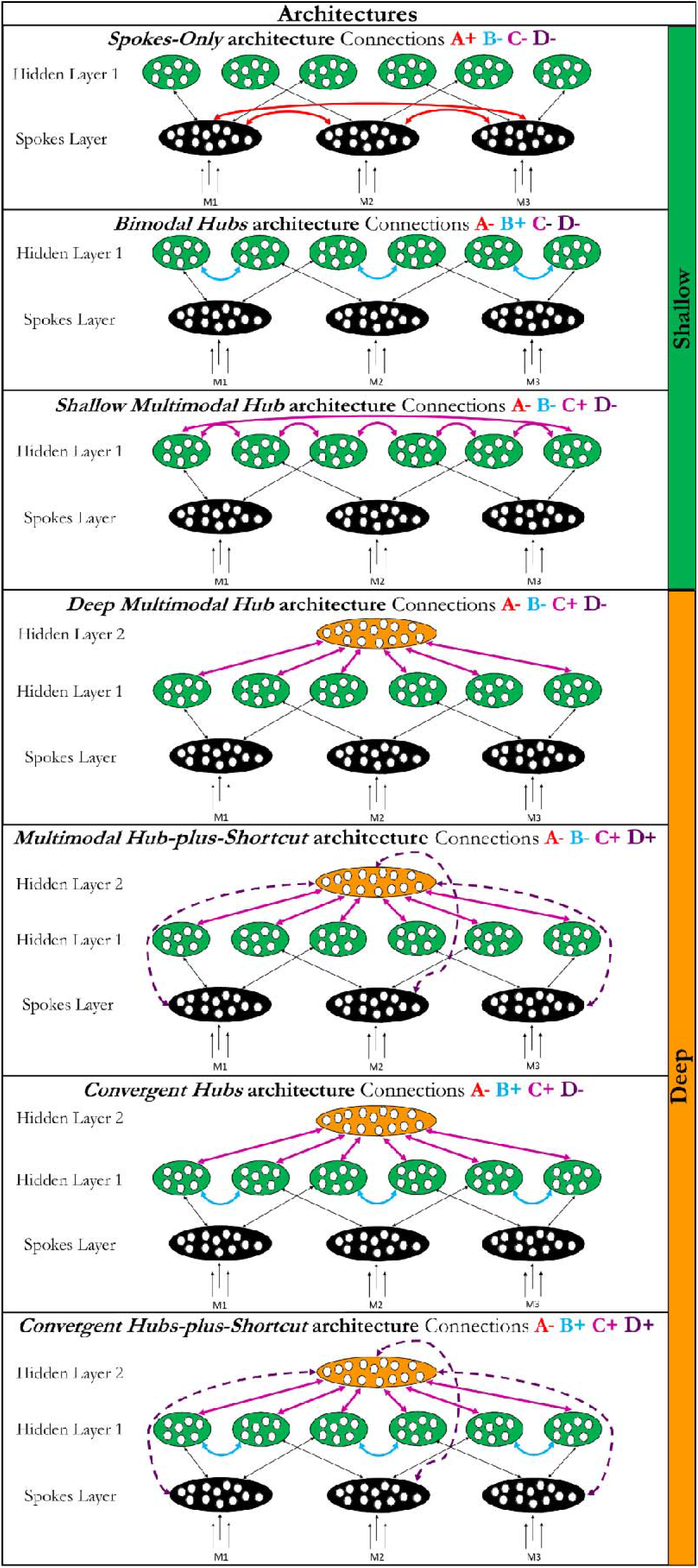
The seven different architectures. Each architecture is based on the shallow or deep configuration and has one or more of the four types of variable connections (A, B, C or D). The presence or absence of each connection type is demonstrated using **+** (where present) or - (where absent). The connection is also shown diagrammatically using arrows in the same colour. Black arrows represent the connections that are stable between architectures. The resulting 7 architectures are provided with labels (italics) for reference within the text. All employ the same total number of weights and units. M = modality.

## Results

The models were fully recurrent continuous-time neural networks trained in the same learning environment to the same performance criterion, differing only in the connectivity amongst units and thus in the organisation of units into layers and pathways (Figure 1). We created activation patterns for 16 items in each of three ‘modalities’ (e.g., word, image, action) designed to capture the central challenge of conceptual abstraction: conceptual structure was latent in the covariance of unit activations across modalities but differed strongly from the covariances apparent within each modality considered independently (Figure 2, see Methods).

**Figure 2.**
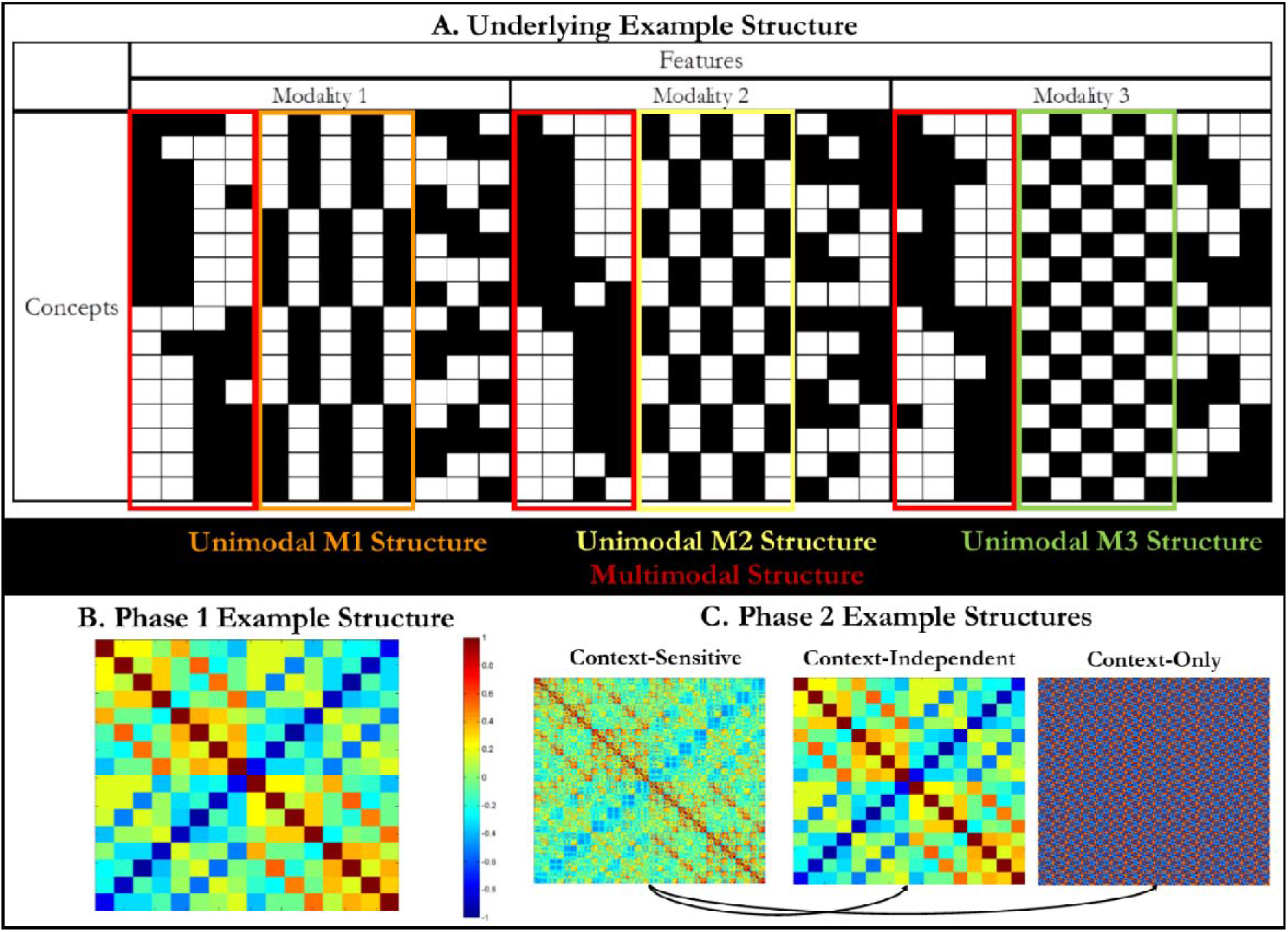
The model environment. A. The full set of features for each concept. Each row is a concept and each column is a feature. A black box indicates that the feature is present and a white box that it is absent for that concept. All 16 concepts are shown here. Red boxes show features that coray strongly across modalities (multimodal structure) whilst the orange, yellow and green boxes highlight features that covary reliably within each modality (unimodal structure). The covariances expressed by the multimodal and each unimodal feature set are mutually orthogonal. B. A matrix showing the context-independent conceptual similarity structure across all modalities for the 16 items. Colours show the correlation (ranging from −1 to 1) for all pairs of vectors concatenated across all columns in Panel A. C. Matrices showing correlations amongst examples used in the context-sensitive simulations in Phase 2, including (left) the full context-sensitive example structure for all 144 input/output patterns, (middle) example structure collapsed across the different task contexts (same as panel B), and (right) similarities amongst the task context representations for each pattern. The 144 patterns arise from crossing 16 items with the 9 possible task contexts. The context-sensitive example structure is a blend of the context-independent conceptual structure used to measure conceptual abstraction (middle) and the context-only similarity structure (right) that indicates the appropriate input and output modalities regardless of concept.

To measure success in conceptual abstraction we computed a ‘true’ conceptual similarity matrix by concatenating the vectors from all three modalities for each item and tabulating the resulting correlations for all item pairs. We used this as the target matrix in a representational similarity analysis^45^: for each model hidden layer, we computed the correlation between the pairwise similarities in its activation patterns and the true conceptual similarities. The highest such correlation for each model indicated its overall success in abstracting conceptual structure (see Supplementary Notes 1 and 2). We considered how this *conceptual abstraction score*, as well as overall speed of learning, varied with model architecture, first without considering context (Phase 1) and then with the additional requirement of context-sensitivity (Phase 2). These simulations strongly favoured one architecture for controlled semantic cognition, which we then assessed in its ability to explain core phenomena in acquired deficits of semantic representation and control (Phase 3).

### Phase 1: Conceptual representation without control

Consistent with prior work^6^, phase 1 models learned to generate, from input provided to any individual modality, the full pattern associated with the concept across all three modalities. The trained models varied dramatically in their conceptual abstraction scores (F(6, 1273)=1168.575, p<.001; Figure 3, Table S1, all significant contrasts had p<.001). Scores were better when the architecture included some form of hub (*Spokes-Only* < *Bimodal Hubs;* t(718)=−38.763) and better still with a multimodal hub *(Bimodal Hubs* < *Shallow Multimodal Hub;* t(718)=−46.634). Depth had no significant effect *(Shallow Multimodal Hub* ~= *Deep Multimodal Hub*; t(105.648)=-0.645, p=.521) and hierarchical convergence had a significant detrimental effect compared to a single multimodal hub *(Deep Multimodal Hub > Convergent Hubs;* t(133.049)=22.503; *Multimodal Hub-plus-Shortcut > Convergent Hubs-plus-Shortcut;* t(114.224)=11.436). Addition of shortcut connections further improved performance for both deep architectures *(Deep Multimodal Hub < Multimodal Hub-plus-Shortcut*; t(138.57)=−5.444; *Convergent Hubs < Convergent Hubs-plus-Shortmt;* t(158)=−14.748). Thus, the *Multimodal Hub-plus-Shortcut* model performed better than all other architectures.

**Figure 3.**
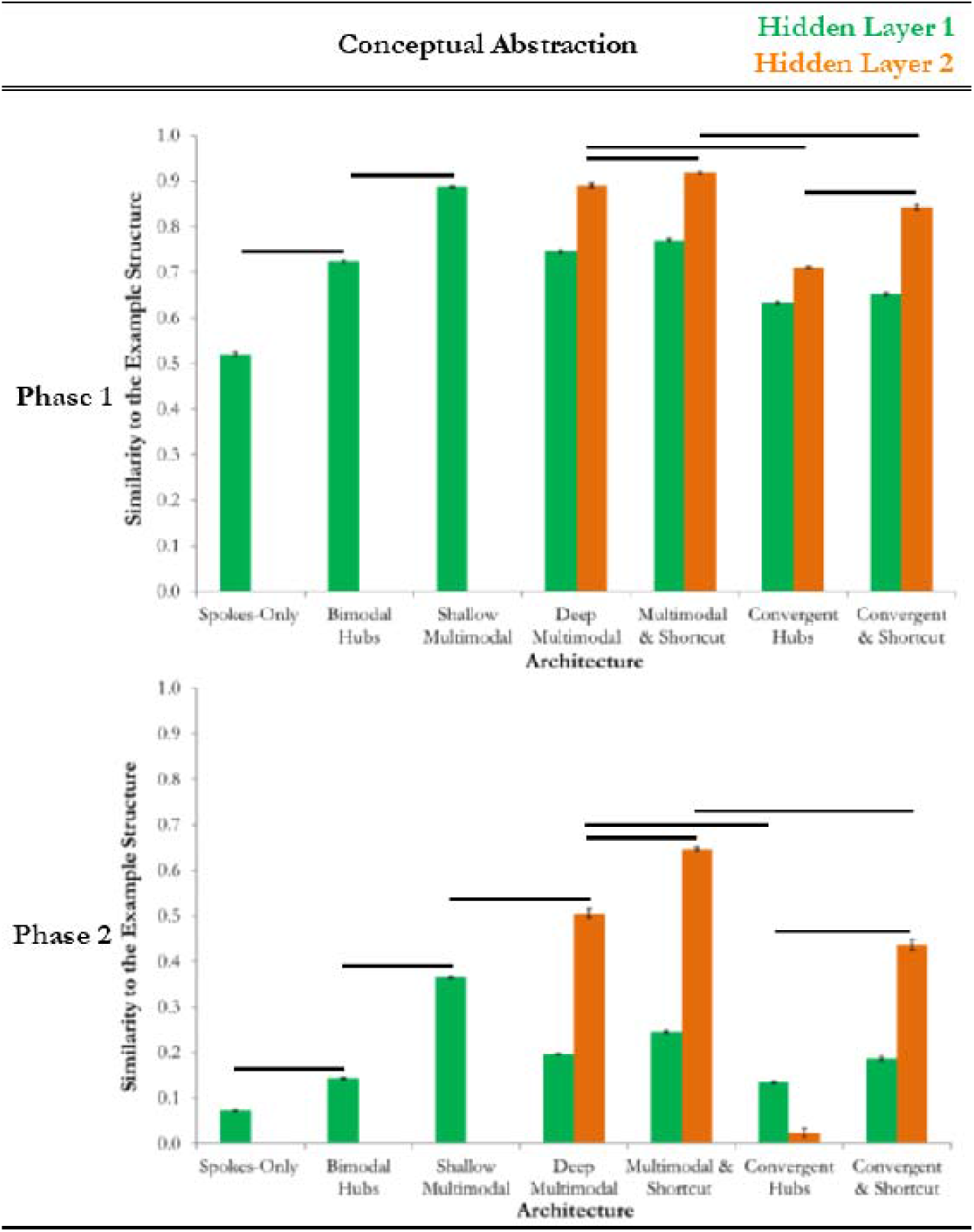
Comparing the conceptual abstraction across the architectures without (in Phase 1) and with (in Phase 2) the additional demand of context-appropriate output. The similarity between the context-independent example structure and the representations in Hidden Layer 1 (green) and Hidden Layer 2 (orange) of each architecture are displayed. The higher bar reflects the conceptual abstraction score for that model architecture. The bar shows the mean similarity value across the different runs of the model (bars show standard error). Planned contrasts with significant differences in the conceptual abstraction score are highlighted with a black line (p<.05).

The number of training epochs required to reach the performance criterion also varied reliably by architecture (F(6, 98)=39.307, p<.001, Figure 4). The *Spokes-Only* structure learned fastest, with the presence of a hub increasing the time taken *(Spokes-Only < Bimodal Hubs*; t(14.163)=−13.057), yet a multimodal hub decreasing it somewhat (*Bimodal Hubs > Shallow Multimodal Hub*; t(15.735)=9.545). Depth slowed training dramatically (*Shallow Multimodal Hub < Deep Multimodal Hub;* t(14.846)=-10.408), but this slowing diminished substantially with shortcut connections (*Deep Multimodal Hub > Multimodal Hub-plus-Shortcut*; t(20.653)=7.152; *Convergent Hubs > Convergent Hubs-plus-Shortcut*; t(15.581)=5.065). Hierarclaical convergence had no effect on learning time *(Deep Multimodal Hub* ~= *Convergent Hubs*; t(28)=−0.622, p=.539; *Multimodal Hub-plus-Shortcut* ~= *Convergent Hubs-plus-Shortcut*; t(28)=-0.213, p=.833).

**Figure 4.**
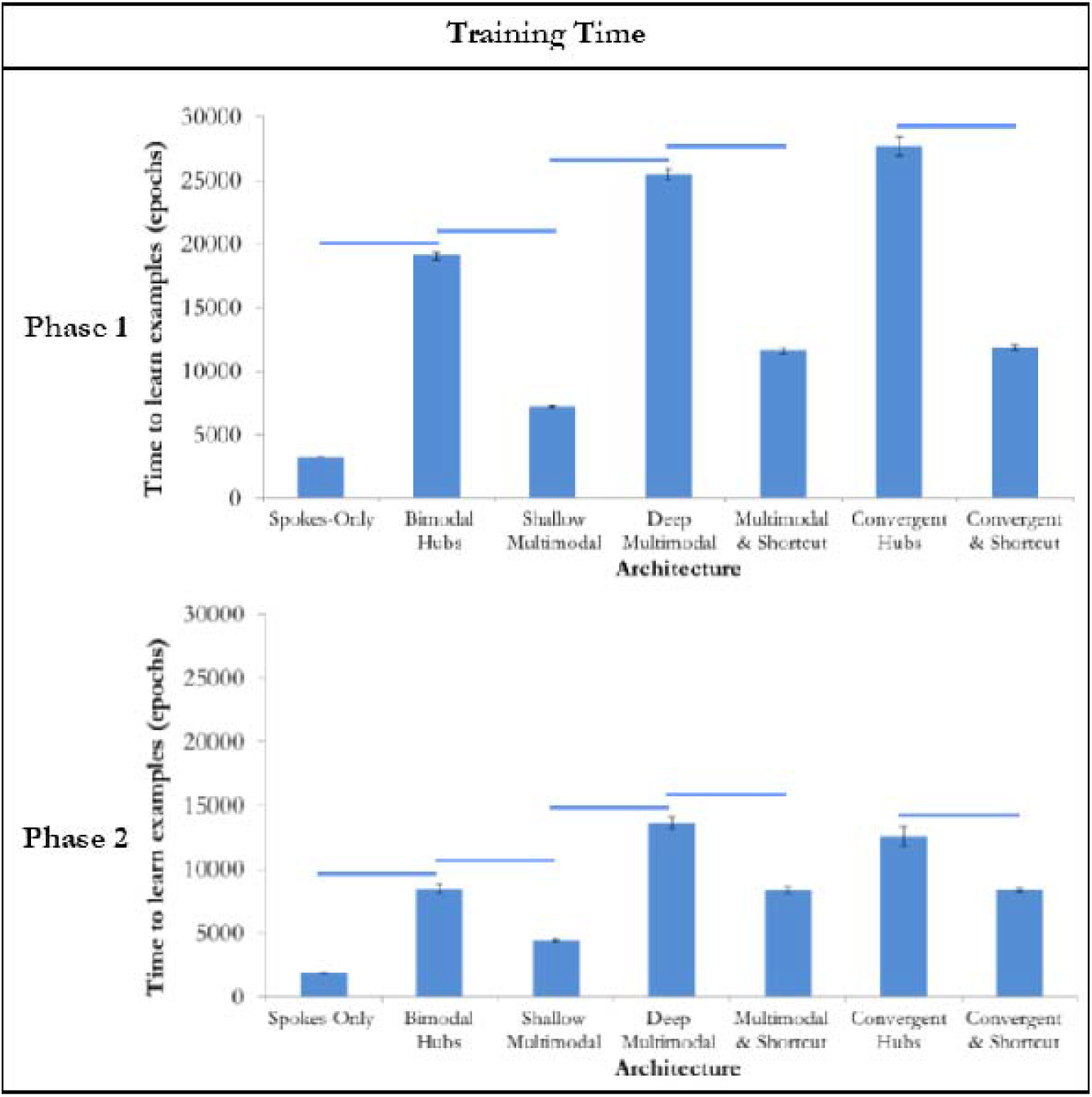
Comparing the training time across the architectures without (in Phase 1) and with (in Phase 2) the additional demand of context-appropriate output. The time taken to learn the examples as the number of epochs of training is displayed. The bars show the mean number of epochs across different runs of the model (bars show standard error). Significant differences in the planned contrasts are highlighted with a line (p<.05).

The efficient abstraction of context-independent conceptual structure depended critically on the presence of a single multimodal hub. Depth alone did not improve the quality of representation and greatly increased training time, but addition of shortcut connections produced the highest-quality representation whilst speeding learning somewhat. Hierarchical convergence reduced representational quality without altering learning time. Of all contrasts, the presence and multimodal nature of a hub produced by far the largest effects. Thus, an interim conclusion from Phase 1 is that conceptual abstraction requires a single multimodal hub.

### Phase 2: Controlled semantic cognition

Phase 2 simulations addressed the full challenge of controlled semantic cognition - performing context-independent conceptual abstraction when experiencing and generating only a limited, context-sensitive subset of an item’s properties in any learning episode. Thus, three additional *context* units were added to each model, each coding the task-relevance of a corresponding input/output modality. For instance, to represent a context in which modality 1 and 2 are important but modality 3 is not (e.g., picture naming, which requires visual input and verbal output without action), context units 1 and 2 would be active while unit 3 would be inactive, and so on. This Control Layer sent trainable unidirectional connections to all units in the network, providing a simple way of implementing control as an influence of the current context on the flow of activation through the network to generate task-appropriate representations and behaviours^6,46,47^ (see Figure 5.A.).

**Figure 5.**
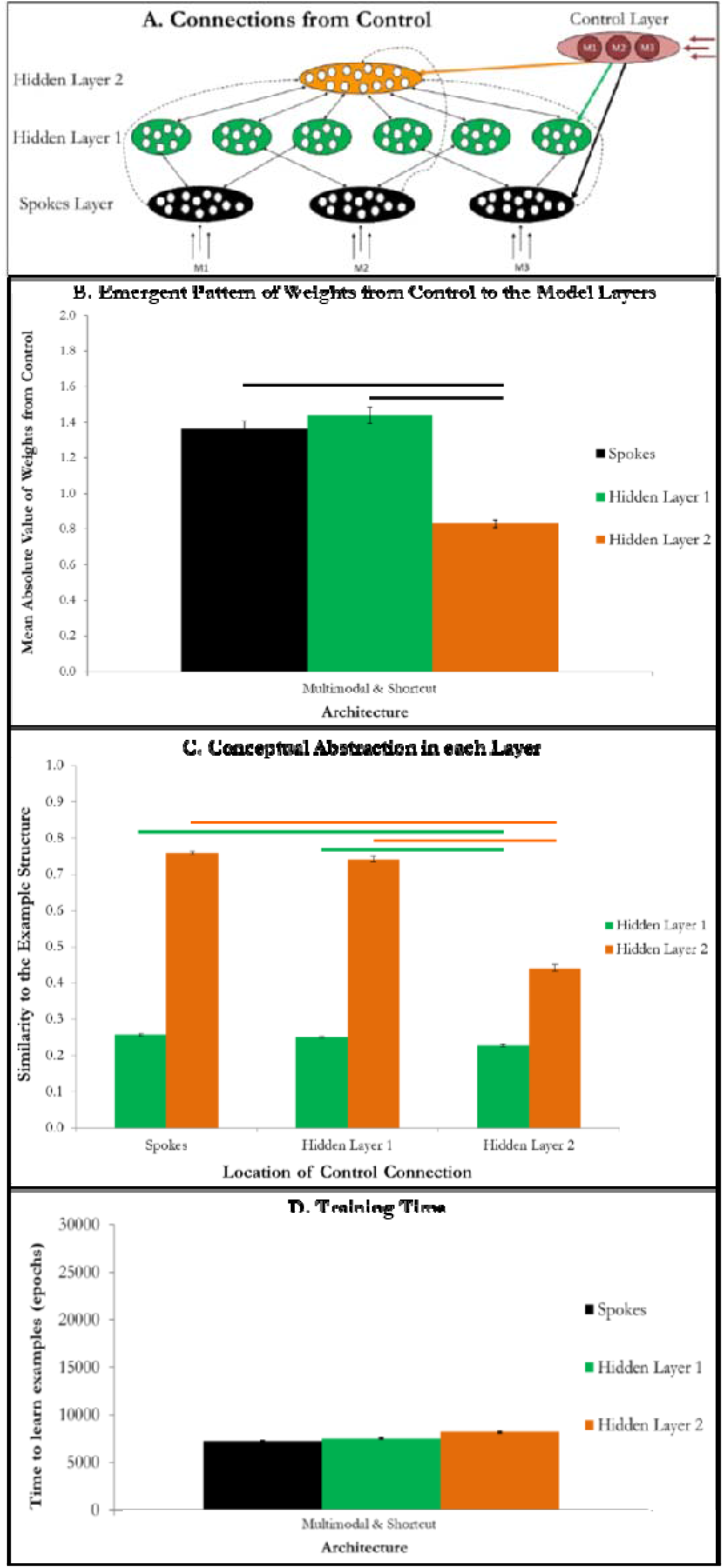
Consequences of the location of the connection to control. A. The different ways control was connected to the Multimodal Hub-plus-Shortcut architecture. The modalities to attend to (i.e., those where an input is received or an output expected) are input to the 3 units in the Control Layer. Learnt unidirectional connections from the Control Layer allow the control signal to enter the semantic system at different points. Connections to all layers were present in initial simulations and the magnitude of weights to each layer compared. Then, the Control Layer was selectively connected to either the Spokes Layer (black arrow), Hidden Layer 1 (green arrow) or Hidden Layer 2 (orange arrow) and the results of these simulations compared. B. The emergent pattern of the weights from the control units to the Spokes Layer, Hidden Layer 1 and Hidden Layer 2 in the Multimodal Hub-plus-Shortcut architecture. The mean absolute value weight to each layer is shown with the standard error. The black bars show significant differences between the weights to the different layers (p<.05). C. The effect of connecting the Control Layer to each layer of the Multimodal Hub-plus-Shortcut architecture on the similarity between the context-independent example structure and the representations in Hidden Layer 1 (green) and Hidden Layer 2 (orange). The bar shows the mean similarity value across the different runs of the model (bars show standard error). The similarity in the deepest layer of an architecture reflects the conceptual abstraction score (orange bar if present, green if not). Significant differences in the planned contrasts are highlighted with a line (p<.05). Significant contrasts in Hidden Layer 1 are shown in green, whereas Hidden Layer 2 differences are shown in orange. D. The time taken to learn the examples is shown as the number of epochs of training. The bars show the mean number of epochs across different runs of the model (bars show standard error) when the control signal is connected to the Spokes Layer (black), Hidden Layer 1 (green) or Hidden Layer 2 (orange). Significant differences in the planned contrasts are highlighted with a line (p<.05).

The models were trained to generate context-sensitive outputs from partial inputs for all 16 items in each of 9 *tasks*, defined by specifying which of the 3 modalities were relevant for input/output. Tasks could involve the same modality for both (e.g., simply viewing an image), or one modality as input and a second as output (e.g., picture naming). From the task representation and an item’s relevant input features, the models learned to activate all task-relevant features for the item while keeping task-irrelevant features inactive. Thus, the models experienced only a limited subset of an item’s features in any given training example in both input and targets, and learned to produce context-sensitive outputs for each item. We then compared the different architectures in their conceptual abstraction ability as in Phase 1.

Success varied substantially by architecture (F(6, 1673)=2326.016, p<.001; Figure 3, Table S2; all significant contrasts had p<.001). Bimodal hubs improved performance (*Spokes-Only < Bimodal Hubs*; t(611.94)=-29.141), but a multimodal hub performed still better (*Bimodal Hubs < Shallow Multimodal Hub*; t(718)=-59.050). In contrast to Phase 1, depth significantly improved conceptual abstraction under conditions of control *(Shallow Multimodal Hub < Deep Multimodal Hub;* t(92.161)=-14.049). Hierarchical convergence dramatically reduced conceptual abstraction (*Deep Multimodal Hub > Convergent Hubs*; t(90.826)=37.123; *Multimodal Hub-plus-Shortcut > Convergent Hubs-plus-Shortcut; t*(137.218)=15.495). Shortcut connections again improved conceptual abstraction in both deep architectures (*Deep Multimodal Hub < Multimodal Hub-plus-Shortcut*; t(148.947)=-l 1.548; *Convergent Hubs < Convergent Hubs-plus-Shortcut*; t(87.551)=-26.027). Only the *Multimodal Hub-plus-Shortcut* architecture acquired representations significantly closer to the context-independent conceptual structure than the control structure (Supplementary Note 3).

Training time also varied by architecture (Figure 4; F(6,98)=113.036, p<.001), with effects mimicking those observed without control. The *Spokes-Only* architecture was fastest, with the addition of a bimodal hub leading to slowing (*Spokes-Only < Bimodal Hubs;* t(14.504)=−15.720), and a multimodal hub reducing this (*Bimodal Hubs > Shallow Multimodal Hub;* t(15.833)=9.424, p<.001). Depth significantly slowed learning (*Shallow Multimodal Hub < Deep Multimodal Hub;* t(15.778)=-21.041, p<.001), yet shortcut connections alleviated this effect *(Deep Multimodal Hub > Multimodal Hub-plus-Shortcut*; t(28)=8.235, p<.001; *Convergent Hubs > Convergent Hubs-plus-Shortcut*; t(28)=6.077, p<.001). Addition of hierarchical convergence did not change learning time *(Deep Multimodal Hub* ~= *Convergent Hubs*; t(28)=1.802, p=.576; *Multimodal Hub-plus-Shortcut* ~= *Convergent Hubs-plus-Shortcut*; t(28)=−0.017, p=1).

In these simulations, context units connected to all units in the semantic network, with their influence shaped by learning. In the best-performing *Multimodal Hub-plus-Shortcut* architecture, learned weights from control to the hub were smaller in magnitude than those projecting to shallower hidden units (t(1517.275)=11.824, p<.001) and spoke units (t(1512.273)=10.364, p<.001) suggesting that control should operate on more superficial layers for the system to acquire context-dependent behaviours (Figure 5.B.). To test this idea, we compared models in which control connected *only* to Spokes, Hidden Layer 1, or Hidden Layer 2 units (Figure 5, Supplementary Note 4). Conceptual abstraction suffered when control operated on the multimodal hub compared to the Spokes layer (t(127.713)=31.981, p<.001) or Hidden Layer 1 only (t(149.191)=27.631, p<.001), which did not differ from one another (p>.1). Moreover, control connectivity to just the spokes (t(138.78)=21.504, p<.001) or shallow hidden units (t(158)=9.493, p<.001) produced reliably better conceptual abstraction than control connecting to all layers, despite the lower number of connections employed; and only these models acquired internal representations significantly more similar to the context-independent conceptual structure than the context structure (Supplementary Note 3). Locus of control did not affect training time (p>.1). Thus, the reverse-engineering approach suggests that controlled semantic cognition, where the system must generate specific responses from partial, context-dependent information, is best achieved within an architecture employing a single, deep multimodal hub and shortcut connections, with control systems acting on superficial rather than deep network components.

### Phase three: Modelling distinct semantic syndromes

Phase 3 tested whether the reverse-engineered model accounts for a different kind of empirical data, namely the distinct patterns of semantic dysfunction arising from damage to representational versus control systems in cortex, as identified in SD and SA patients, respectively. Whilst a detailed accounting of these differences is beyond the scope of this paper, prior research has revealed similarities and differences between syndromes. Both can show comparably severe semantic deficits with frequent omissions in various tasks, but they differ in errors of commission. Patients with SA more often generate task-inappropriate responses, like producing associative errors (“acorn” for squirrel) and circumlocutions (“has stripes” for zebra) in naming, losing track of the target category in verbal fluency (e.g., for birds: robin, sparrow, chicken, pig, cow), or failing to grasp a tool in a manner that affords its correct use in a given task context^2,13,48^. In contrast, patients with SD more often generate task-appropriate but semantically incorrect behaviours: committing semantic (e.g., “horse” for zebra) or ordinacy (e.g., “animal” for squirrel) errors in naming, generating fewer but mainly correct items in verbal fluency, and grasping a tool correctly but exhibiting a semantically inappropriate use (e.g., brushing hair with a comb)^2,10,13,49^.

In the optimal model from Phase 2, we simulated disordered control in SA by adding noise to the control units activations, and degraded representation in SD by removing a proportion of connections to, from and within the multimodal hub ^30^. We then assessed performance under increasing levels of simulated damage for each model syndrome, matched for severity as indexed by the total number of errors (Figure 6). From these data we compared the relative frequency of three error types across the two forms of damage: omissions (inactivation of a correct features), context-appropriate commissions (activation of an incorrect but task-relevant feature), and context-inappropriate commissions (activation of a task-irrelevant feature).

**Figure 6.**
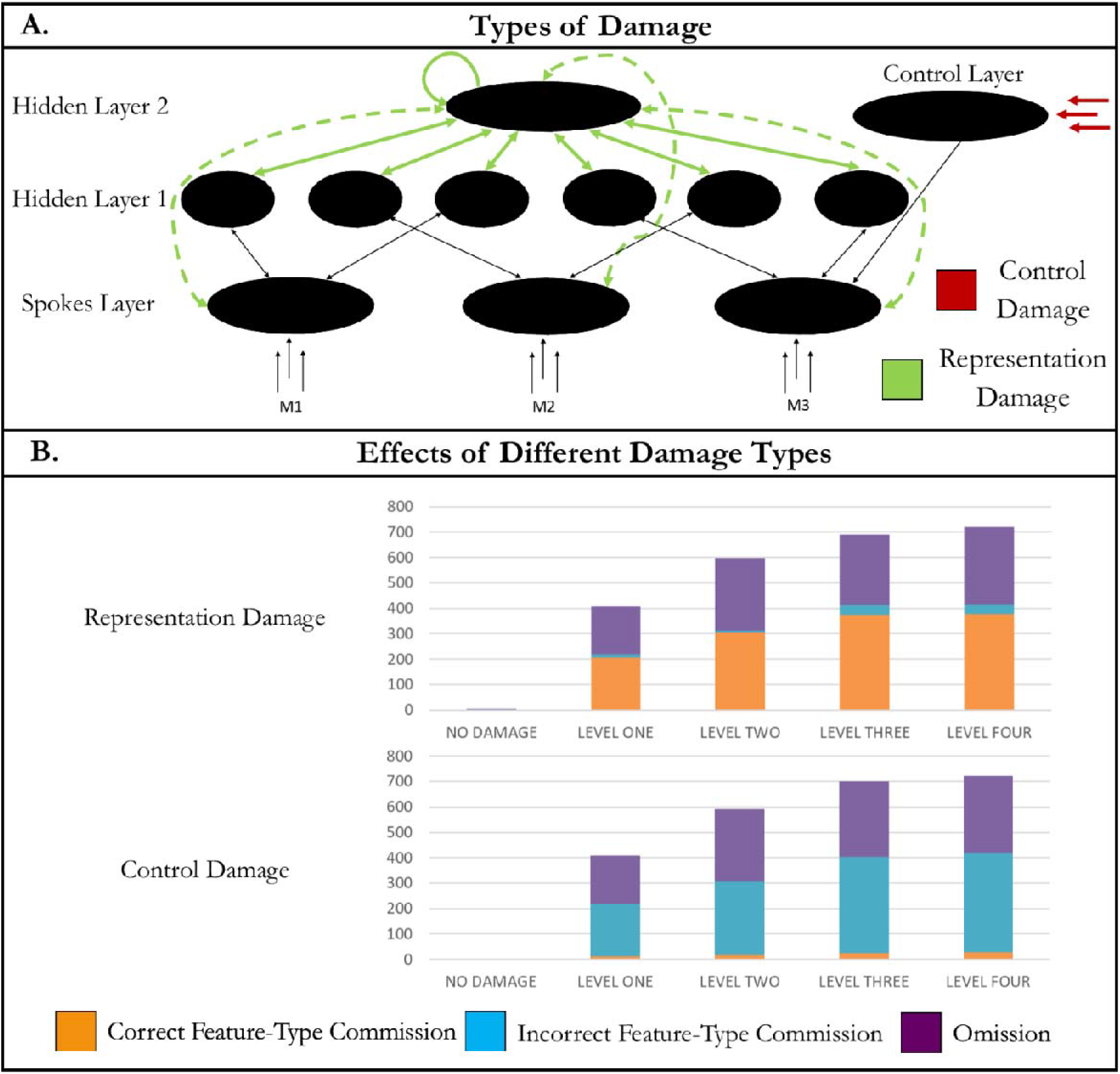
The effects of damage to control and representation within the Multimodal Hub-plus-Shartcut architecture. A. The connections and inputs affected in the different patterns of damage. The Representation Damage simulation involved removing a proportion of all connections within, to and from Hidden Layer 2 (shown in green). The Control Damage simulation involved adding noise to the input to the control units (shown in red). B. The different kinds of feature errors resulting from each type of damage. The height of the bar shows the number of errorful features, which is matched across damage types at each level. Each errorful feature is an omission of a correct features (purple), a commission of a feature of the correct type (orange), or a commission of a feature of the incorrect type (blue). The two damage types have qualitatively different patterns.

At every level, damage to control produced fewer context-appropriate commissions (damage type; F(1, 190)=1292.758, p<.001; damage level; F(4,190)=128.784, p<.001; interaction; F(4,190)=99.301, p<.001) and more context-inappropriate commissions (damage type; F(1, 190)=2194.628; damage level; F(4,190)=168.893, p<.001; interaction; F(4,190)=168.893, p<.001) than damage to representation (Supplementary Note 5). Feature omissions were equally frequent and reflected damage severity (damage type; F(1,190)=0.613, p=.435; damage level; F(4,190)=440.445, p<.001; interaction; F(4,190)=0.570, p=0.685). Thus, the reverse-engineered architecture accounts for qualitatively different patterns of impaired semantic cognition arising from damage to control versus representational elements of the system.

## Discussion

We applied a novel reverse-engineering approach to discover a neural network architecture capable of achieving the core, opposing functions of controlled semantic cognition: conceptual abstraction across modalities and contexts with simultaneous context-sensitivity. The optimal network had four important characteristics: (1) a single multimodal hub, (2) a deep architecture, (3) sparse shortcut connections, and (4) control operating on shallow rather than deep network components. The reverse-engineered model additionally accounted for similarities and differences in semantic impairment arising from damage to systems of representation vs. control. In this discussion we consider why these elements are critical, how the reverse-engineered architecture relates to the known anatomy of the cortical semantic system, and implications of the work for theories of controlled semantic cognition.

### Why a single multimodal hub?

Prior work established that feedforward neural networks exploit shared structure across modalities and contexts only when information from each gets passed through the same units and weights somewhere in the network, an observation termed the *convergence principle*^6,30^. Weights that contribute to processing for all items, modalities, and contexts get shaped by all learning episodes, providing a medium for encoding shared structure. It was unclear whether the same principle operates in recurrent networks where all weights and units potentially contribute to processing for all varieties of information. This question is important because most neuro-semantic models assume recurrent, interactive processing. Here we establish that convergence remains critical for learning structure in recurrent networks: architectures lacking a single multimodal hub can learn the same input/output mappings, but do not acquire internal representations that reflect the full conceptual representational structure across modalities and learning episodes. Even models possessing a multimodal hub fail to learn the desired structure if they also possess shallower and more direct pathways between modalities. Here the mappings between modalities can be acquired via the shallower pathways, with little pressure to use the connections mediating all modalities. Only architectures that constrain the network to use these weights acquire the important conceptual structure. These findings are problematic for distributed-only^9,50^ and multi-hub^26,27^ theories of semantic cognition, yet consistent with the Hub-and-Spoke theory^30^. The current simulations provide a computational instantiation of the Hub-and-Spoke and Controlled Semantic Cognition frameworks^1,30^ with additional computational constraints; depth (also instantiated in^40^) and shortcut connections.

### Why deep representation and shallow control?

Deep networks can acquire complex internal representations that generalise well when trained on large corpora of naturally occurring stimuli^51^. Yet in the current work, deep and shallow networks possessing a multimodal hub learned the target structure equally well when trained without context. Only when required to generate context-sensitive outputs did the deeper model outperform the shallow model. Thus, depth may have a particular benefit for controlled semantic cognition; specifically facilitating the network’s ability to discover representational structure across modalities and context when learning involves experience with limited, context-dependent inputs and outputs.

Why does depth help? Different tasks activate different surface properties for the same item: to retrieve an item’s colour the system must activate visual and not action properties, whilst the reverse holds when retrieving the item’s function. Moreover, the inter-item similarities expressed in different modalities can differ dramatically e.g., a fire-extinguisher and ketchup share colour but differ radically in function. Context-sensitive training thus pressures the system to represent a given item quite differently in different task contexts, making it difficult for the system to detect covariance of properties across contexts. Feedforward models can overcome such pressure when their architecture ensures that contextual information does not directly influence conceptual representations^6^. The current work shows a similar phenomenon in the recurrent case when the network is deep and control operates on its shallower elements. If the multimodal hub connects directly to unimodal representations (i.e., the model is shallow), context strongly influences the representations it acquires. Likewise, a deep model in which the multimodal hub directly receives context inputs acquires context-bound representations. Only when the model is deep and control operates on the shallower elements is the hub sufficiently insulated from contextual information to acquire more context-invariant representations that express cross-context structure.

### Why shortcut connections?

Deep networks initially learn extremely slowly due to the “vanishing gradient” problem^52^: with many layers of weights intervening between input and output, changes to earlier weights produce negligible impact on output patterns, so error-driven learning produces minimal weight changes. Likewise, different input patterns initially elicit almost undifferentiated activation patterns in deeper network layers, making it difficult for downstream weights to produce different outputs for different stimuli. In fully-connected deep networks these factors produce an initial fallow period with little reduction in error^6,53^. The current results show that shortcut connections can significantly remediate this problem. Activations and error signals propagate through fewer layers in the shortcut pathways so these weights learn more quickly. These shortcut weights drive better-differentiated activation patterns in the middle and output layers of the network, boosting the propagation of error derivatives throughout the deeper, fully-connected pathway and speeding learning overall. Such shortcut connections can produce a remarkable increase in speed of learning even when very sparse.

### Accounting for distinct semantic syndromes

The simulations showed that damage to control versus hub representations can produce equivalently severe impairment with qualitative differences in error types: control damage elicited more errors for context-inappropriate features whereas representation damage elicited more errors for context-appropriate features. These distinct feature error types explain the differential pattern of item-level errors in SA and SD. In the control deficit only, the inclusion of additional context-inappropriate features would lead to contextually-inappropriate behaviour, whereas with representation damage errors only occur within a context, resulting in task-appropriate but semantically incorrect behaviours. Why do these differential error types occur?

In the intact model, control can be viewed as “selecting” which properties matter for the task; potentiating context-relevant surface properties (making them easier to activate) while suppressing context-irrelevant properties (making them harder to activate^47^). Correct context-sensitive responding arises from the joint influence of representational and control systems on surface properties. A control signal distorted by noise will incorrectly potentiate some context-irrelevant units while suppressing some context-relevant units. In this case input from the intact representational network will activate some task-irrelevant units and fail to activate some item-and context-appropriate units. Conversely when damage affects the hub representation, the control signal remains intact—only context-appropriate features are potentiated and therefore only context-appropriate commissions can occur.

### Connection to established neuroanatomy

The reverse-engineered architecture mirrors the known anatomy of the cortical semantic system remarkably well. The ventral ATL forms a multimodal conceptual hub, as demonstrated in SD^10-12^, brain imaging^32,54-56^, neurostimulation^36,37^ and intracortical electrode recording^34,57^. The progression from unimodal perceptual representations to multimodal conceptual representations occurs in a graded fashion across many cortical areas^55,58^ corresponding well to a deep network. Long-distance white matter tracts, such as the inferior longitudinal fasciculus^59^, traverse temporal cortex, providing pathways for sparse shortcut connectivity. Recent evidence suggests specifically that posterior fusiform connects to the ATL both via gradual progression along the anterior fusiform and a ‘direct’ white matter route^60^.

The model also captures the functional and anatomical division between systems of semantic representation in anterior temporal regions vs. systems of semantic control supported by fronto-parietal cortex. Several lines of evidence suggest that these cortical regions serve different functions in semantic cognition, including (1) the distinct semantic impairments arising from damage to anterior temporal *vs.* frontoparietal cortex already discussed^2,13^, (2) the degree to which these regions show context-dependent activation^2,61,62^, and (3) differing patterns of connectivity for each region^63,64^. The current simulations elucidate why the cortical semantic network is structured this way. The anatomy of the temporal lobe promotes the extraction of conceptual structure across modalities and time in the anterior multimodal hub, whilst still allowing for context-dependent activation patterns within unimodal “spoke” regions. This relative functional specialisation, alongside the anatomical separation of representation and control systems promotes discovery of context-independent representations in tandem with context-appropriate behaviour.

Finally, the reverse-engineered model suggests that semantic control should exert more direct influence over shallower elements of the representational network. Classic functional imaging studies support this hypothesis. For instance, retrieval of colour information from a word stimulus preferentially activates cortical areas near those that support colour perception, whilst retrieval of action information from the same stimulus preferentially activates regions near those supporting motion perception^65^. These and related studies^66^ suggest that task context representations modulate the activation of content-specific representations within the spokes of the semantic network. Precisely how control and representation regions are structurally connected remains an open question. The current simulations provide a testable hypothesis: control regions should not structurally connect directly to the anterior temporal hub. Whilst the literature does not definitively answer this question, the core ventral ATL hub region does have few connections to distal regions^67,68^. Further work is needed to extend our understanding of the controlled semantic network and delineate its precise cortical loci.

## Method

### Model Environment & Control

Each concept consisted of 12 features in each of 3 modalities (M1, M2 and M3; see Figure 2). Concepts were constructed based on a critical aspect of conceptual structure; unimodal perceptual structures only weakly correlate with the conceptual structure which is more predictive but requires extraction across modalities. The model environment included four orthogonal structures; one distinct unimodal (based on 5 perfectly correlated or anti-correlated features within a single modality) structure per modality (unimodal M1, unimodal M2 and unimodal M3) and a multimodal (based on 12 highly correlated features spread across all three modalities) structure. In each modality the unimodal structure is greater, yet overall the multimodal structure is stronger. Whilst the main analyses focus on the full structure, highly consistent results are displayed for the unimodal and multimodal structures in Supplementary Notes 6 and 7. Input was always in a single modality, with the other two modalities set to 0. For simulations without control, the target was the full concept, resulting in 48 versions of the 16 examples. For simulations with control, each concept was presented in one of three modalities with a control signal designating a required output in one of three modalities (as well as the input modality), resulting in 144 versions of the 16 examples. Task-irrelevant modalities had targets of 0.

### Model Architecture

Code for replicating all simulations is available in the Supplementary Materials. All architectures utilised a single framework, consisting of 12 pairs of input and output units per modality (connected on a one-to-one basis with a frozen weight of 6 and a fixed bias of −3 for the output units), 60 hidden units and 3132 bidirectional connections with learnable weights. Matching the number of resources allowed clear interpretation of the differences between the architectures. All architectures had connections between the three modality-specific regions of the Spokes Layer and the six subsections of Hidden Layer 1 (with two subsections connected to each modality-specific spoke region) and within each portion of Hidden Layer 1. Deep architectures had connections from Hidden layer 1 to Hidden Layer 2 and within Hidden Layer 2. To match the number of connections between architectures, some connections were sparse (see Supplementary Method 1). Two factors varied between the 7 model architectures; the hidden layer configuration (shallow; a single layer of 60 units *vs.* deep; one layer of 42 units and a deeper layer of 18 units) and the presence or absence of four types of connections (A; connections between modality-specific output units in the Spokes Layer; B; connections between pairs of Hidden Layer 1 regions that receive different modalities of input, resulting in the formation of bimodal hubs; C; connections between hidden units to form a single multimodal hub, either within Hidden Layer 1 or Hidden Layer 2; D; direct but sparse shortcut connections between the Spokes Layer and Hidden Layer 2 that bypass Hidden Layer 1). Whilst long-range connections are likely to be relatively sparse, their precise sparsity is not known. D connections were included at a sparse but non-trivial proportion of 1 in 24 (although see Supplementary Note 8 for an assessment of systematically varying the sparsity within the *Multimodal Hub-plus-Shortcut* architecture).

Figure 1 represents each architecture. Three architectures were constructed from the shallow configuration; a ‘*Spokes-Only’* architecture employing A connections only, a *‘Bimodal Hubs’* architecture with B connections only and a *‘Shallow Multimodal Hub’* architecture with C connections only, resulting in Hidden Layer 1 forming a single multimodal hub. All four deep architectures have C connections resulting in a multimodal hub in Hidden Layer 2. The *‘Deep Multimodal Hub’* architecture has no additional connections, thus three modality-specific routes connect via a deep multimodal hub. The *‘Multimodal Hub-plus-Shortcut’* architecture also included D connections and the *‘Convergent Hubs’* architecture included additional B connections, resulting in hierarchical convergence as multiple bimodal hubs connect to a single deep multimodal hub. The *‘Convergent Hubs-plus-Shortcut* architecture combined the B, C and D. The seven architectures allowed contrasts separating the effect of each architectural feature; the effects of a hub (*Spokes-Only vs. Bimodal Hubs)*, a multimodal hub (*Bimodal Hubs vs. Shallow Multimodal Hub)*, depth (*Shallow Multimodal Hub vs. Deep Multimodal Hub)*, shortcut connections *(Deep Multimodal Hub vs. Multimodal Hub-plus-Shortcut* and *Convergent Hubs vs. Convergent Hubs-plus-Shortcut)* and hierarchical convergence *(Deep Multimodal Hub vs. Convergent Hubs* and *Multimodal Hub-plus-Shortcut vs. Convergent Hubs-plus-Shortcut).*

In Phase 2, a ‘Control Layer’ consisting of three units (each corresponding to one modality) was added to provide a context signal. The models had unidirectional learnable connections from the control units to the Spokes Layer, Hidden Layer 1 and Hidden Layer 2 (where present). Initially, no assumptions were made as to where control should connect, allowing a fair comparison across architectures. Following this analysis, the emergent reliance on the connections to each layer was investigated using an equal number of connections to all layers (81 per layer if shallow, 54 per layer if deep). Then, the effectiveness of this emergent pattern was verified by contrasting versions of the model where the Control Layer was connected to each single layer (with the same number of connections).

### Training Parameters

The models were constructed and trained using the Light Efficient Network Simulator (LENS, version 2.63) software ^69^. Each simulation employed a fully recurrent network with 24 activity updates per example (6 time intervals and 4 ticks per time interval). Inputs were presented for the first 3 time intervals. Each training batch consisted of all examples presented once in a random order. At the end of each batch, error derivatives were calculated and all weights in the model adjusted by a small amount. All simulations employed the same training parameters, found to allow learning in pilot simulations. The models were trained using gradient descent with a learning rate of 0.001 and a weight decay parameter of 0.0001 with no momentum. Training ended when all output feature units were within 0.2 of their target. Thus, all architectures were matched on accuracy. Analyses were performed using the final time step of a test trial. Each simulation was performed 60 times.

### Assessment Metrics

Data processing was performed in MATLAB and statistics in the Statistical Package for the Social Science (SPSS, 2013). The similarity structure of the models representations were compared to the ground-truth similarity structure to determine each architectures ability to accurately discover and represent the full structure in the environment. The critical example structure used to form this conceptual abstraction score is the *context-independent* semantic representation structure (the relationships between examples based on the full set of features regardless of the current input or output domain). For the simulations with control it is also possible to look at the similarity of the representations to the context signal (*context-only*) or the full structure varying by context and concept (*context-sensitive*), see Supplementary Note 9.

Correlation-based similarity matrices were constructed from the activity in a model region across all examples after learning. Model regions were defined as portions of the model with the same potential connections (before sparsity is taken in to account) as connectivity constrains function^43,71,72^. This resulted in 3 Hidden Layer 1 regions in the *Spokes-Only, Shallow Multimodal Hub, Deep Multimodal Hub* and *Multimodal Hub-plus-Shortcut* architectures and 6 in the *Bimodal Hubs, Convergent Hubs* and *Convergent Hubs-plus-Shortcut* architectures. The similarity between each result-based similarity matrix and the example-based similarity matrix was determined using a correlation. This resulted in a value per model run and layer subregions for statistical comparisons, although these equivalent values are averaged when reported. The values for the region with the highest similarity to the *context-independent* semantic representation were used to contrast the models (although for comparison of all regions and consideration of the effect of the number of units see Supplementary Note 1). For the simulations with and without control, a repeated measures ANOVA assessed the differences between the 7 architectures and *a priori* between-samples t-tests were used to compare the effect of each architectural feature with Bonferroni correction for the seven multiple comparisons. The number of epochs taken to train each architecture to criterion was determined for 15 runs of each model.

To determine how the Control Layer should connect to the rest of the model, two assessments were used. Firstly, 40 models were ran with connectivity to each layer. The emergent preference for receiving and employing the control signal in each layer was examined by contrasting the sum absolute magnitude of the weights to each layer using 3 t-tests (Spokes vs. Hidden Layer 1, Spokes vs. Hidden Layer 2, Hidden Layer 1 vs. Hidden Layer 2) in the deep architectures and one (Spokes vs. Hidden Layer 1) in the shallow architectures. Bonferroni multiple comparison correction for 3 contrasts was applied to the deep architectures. Secondly, the effectiveness of this emergent pattern was verified by only connecting the models to one layer (either the Spokes Layer, Hidden Layer 1 or, where possible, Hidden Layer 2). These model versions were compared on their extraction of the context-independent representation structure using between-samples t-tests and Bonferroni correction applied.

### Lesioning the Model

To assess the effect of lesions to representation and control regions, the models were constructed and trained using the optimal architecture identified within Phase 2 (including connections from control to the Spokes Layer only). To damage representational processes, the connections to, from and within Hidden Layer 2 were removed as this region had the greatest conceptual abstraction score and thus, showed the greatest specialisation for representation processes. In the Control Damage simulations, noise was added to the input to the control units. Three types of errors may be made; omission of a feature that is correct both for that concept and that context, commission of a feature that is in the correct context but incorrect for that concept and commission of a feature in the incorrect context. To allow comparison across damage type, controlling for the effect of damage severity, each simulation was performed at a variety of levels with the proportion of weights removed (for Representation Damage) or the amount of noise added (for Control Damage simulations) varied systematically. Then, points at which the number of errorful features (those further than 0.2 from the correct output) were matched across the damage types were identified. At the chosen levels, t-tests showed the three damage types did not have significantly different numbers of errors (each p>.25). This resulted in the identification of four damage levels at which the effect of damage type on the three possible error types could be assessed; Representation Damage with the removal of connections at proportions of 0, 0.1, 0.25, 0.3 and 0.35, and Control Damage with noise added to the control signal with ranges of 0, 0.625, 1, 1.25 and 1.375. For each error type (Correct Feature-Type Commission, Incorrect Feature-Type Commission, Omission) an ANOVA was performed to assess the effects of damage type (Representation Damage, Control simulations) and damage level (No damage, Level 1, Level 2, Level 3, Level 4). Error types were compared across the damage types using independent samples t-tests at each level.

## Acknowledgements

This work was supported by a British Academy Postdoctoral Fellowship awarded to RLJ (pf170068), a programme grant to MALR and TTR from the Medical Research Council (grant number MR/R023883/1), and an Advanced Grant from the European Research Council to MALR (GAP: 670428).

## Author Contributions

RJ, TTR and MLR made substantial contributions to the conception and design of the work, the interpretation of data and the manuscript revisions. RJ acquired the results and drafted the manuscript.

## References

1 Lambon Ralph, M. A., Jefferies, E., Patterson, K. & Rogers, T. T. The neural and computational bases of semantic cognition. Nat. Rev. Neurosci. 18, 42–55 (2017).

2 Jefferies, E. The neural basis of semantic cognition: Converging evidence from neuropsychology, neuroimaging and TMS. Cortex 49, 611–625, doi:10.1016/j.cortex.2012.10.008 (2013).

3 Abel, T. J. et al. Direct physiologic evidence of a heteromodal convergence region for proper naming in human left anterior temporal lobe. J. Neurosci. 35, 1513–1520 (2015).

4 Wittgenstein, L. Philosophical Investigations. (Blackwell Publishing, 1953).

5 Lambon Ralph, M. A., Sage, K., Jones, R. W. & Mayberry, E. J. Coherent concepts are computed in the anterior temporal lobes. Proc. Natl. Acad. Sci. U. S. A. 107, 2717–2722 (2010).

6 Rogers, T. T. & McClelland, J. L. Semantic cognition: A parallel distributed processing approach. (MIT Press, 2004).

7 Saffran, E. M. The organization of semantic memory: In support of a distributed model. Brain Lang. 71, 204–212 (2000).

8 Thompson-Schill, S. L., D’Esposito, M., Aguirre, G. K. & Farah, M. J. Role of left inferior prefrontal cortex in retrieval of semantic knowledge: A reevaluation. Proc. Natl. Acad. Sci. U. S. A. 94, 14792–14797 (1997).

9 Eggert, G. H. Wernicke’s works on aphasia: A sourcebook and review. (Mouton, 1977).

10 Patterson, K., Nestor, P. J. & Rogers, T. T. Where do you know what you know? The representation of semantic knowledge in the human brain. Nat. Rev. Neurosci. 8, 976–987, doi:10.1038/nrn2277 (2007).

11 Acosta-Cabronero, J. et al. Atrophy, hypometabolism and white matter abnormalities in semantic dementia tell a coherent story. Brain 134, 2025–2035 (2011).

12 Warrington, E. K. Selective impairment of semantic memory. Quarterly Journal of Experimental Psychology 27, 635–657, doi:10.1080/14640747508400525 (1975).

13 Jefferies, E. & Lambon Ralph, M. A. Semantic impairment in stroke aphasia versus semantic dementia: a case-series comparison. Brain 129, 2132–2147, doi:10.1093/brain/awl153 (2006).

14 Rosch, E., Mervis, C. B., Gray, W., Johnson, D. & Boyes-Braem, P. Basic objects in natural categories. Cogn. Psychol. 8, 382–439 (1976).

15 Murphy, G. L. & Medin, D. L. The role of theories in conceptual coherence. Psychol. Rev. 92, 289–316 (1985).

16 Keil, F. C. in The Jean Piaget Symposium series. The epigenesis of mind: Essays on biology and cognition (eds S. Carey & R. Gelman) 237–256 (Lawrence Erlbaum Associates, Inc., 1991).

17 Barsalou, L. W. Perceptual symbol systems. Behavioral and Brain Sciences 22, 577–660 (1999).

18 Gelman, S. A., Leslie, S. J., Was, A. M. & Koch, C. M. Children’s interpretations of general quantifiers, specific quantifiers and generics. Language, Cognition and Neuroscience 30, 448–461 (2015).

19 Rumelhart, D. E. & Todd, P. M. in Attention and performance XIV: Synergies in experimental psychology, artificial intelligence, and cognitive neuroscience (eds D. E. Meyer & S. Kornblum) 3–30 (MIT Press, 1993).

20 Martin, A. & Chao, L. L. Semantic memory and the brain: structure and processes. Current Opinion in Neurobiology 11, 194–201 (2001).

21 Huth, A. G., de Heer, W. A., Griffiths, T. L., Theunissen, F. E. & Gallant, J. L. Natural speech reveals the semantic maps that tile human cerebral cortex. Nature 532, 453–458 (2016).

22 McCrae, K., de Sa, V. R. & Seidenberg, M. S. On the nature and scope of featural representations of word meaning. Journal of Experimental Psychology: General 126, 99–130 (1997).

23 Lambon Ralph, M. A., McClelland, J. L., Patterson, K., Galton, C. J. & Hodges, J. R. No right to speak? The relationship between object naming and semantic impairment: Neuropsychological abstract evidence and a computational model. J. Cogn. Neurosci. 13, 341–356 (2001).

24 Farah, M. J. & McClelland, J. L. A computational model of semantic memory impairment: Modality specificity and emergent category specificity. Journal of Experimental Psychology: General 120, 339–357 (1991).

25 Devereux, B. J., Clarke, A. & Tyler, L. K. Integrated deep visual and semantic attractor neural networks predict fMRI pattern-information along the ventral object processing pathway. Scientific Reports 8, 10636 (2018).

26 Binder, J. R. & Desai, R. H. The neurobiology of semantic memory. Trends Cogn. Sci. 15, 527–536, doi:10.1016/j.tics.2011.10.001 (2011).

27 Damasio, H., Grabowski, T. J., Tranel, D., Hichwa, R. D. & Damasio, A. R. A neural basis for lexical retrieval. Nature 380, 499–505 (1996).

28 Damasio, A. R. & Damasio, H. in Computational neuroscience. Large-scale neuronal theories of the brain (eds C. Koch & J.L. Davis) 61–74 (MIT Press, 1994).

29 Mahon, B. Z. & Caramazza, A. A critical look at the embodied cognition hypothesis and a new proposal for grounding conceptual content. J. Physiol.-Paris 102, 59–70 (2008).

30 Rogers, T. T. et al. Structure and deterioration of semantic memory: A neuropsychological and computational investigation. Psychol. Rev. 111, 205–235, doi:10.1037/0033-295x.111.1.205 (2004).

31 Lambon Ralph, M. A., Lowe, C. & Rogers, T. T. Neural basis of category-specific semantic deficits for living things: evidence from semantic dementia, HSVE and a neural network model. Brain 130, 1127–1137 (2007).

32 Binney, R. J., Embleton, K. V., Jefferies, E., Parker, G. J. M. & Lambon Ralph, M. A. The ventral and inferolateral aspects of the anterior temporal lobe are crucial in semantic memory: Evidence from a novel direct comparison of distortion-corrected fMRI, rTMS, and semantic dementia. Cereb. Cortex 20, 2728–2738, doi:10.1093/cercor/bhq019 (2010).

33 Visser, M., Jefferies, E., Embleton, K. V. & Lambon Ralph, M. A. Both the middle temporal gyrus and the ventral anterior temporal area are crucial for multimodal semantic processing: Distortion-corrected fMRI evidence for a double gradient of information convergence in the temporal lobes. J. Cogn. Neurosci. 24, 1766–1778 (2012).

34 Shimotake, A. et al. Direct exploration of the ventral anterior temporal lobe in semantic memory: Cortical stimulation and local field potential evidence from subdural grid electrodes. Cereb. Cortex (2014).

35 Matsumoto, R. et al. Functional connectivity in the human language system: a cortico-cortical evoked potential study. Brain 127, 2316–2330, doi:10.1093/brain/awh246 (2004).

36 Pobric, G., Jefferies, E. & Lambon Ralph, M. A. Anterior temporal lobes mediate semantic representation: Mimicking semantic dementia by using rTMS in normal participants. Proc. Natl. Acad. Sci. U. S. A. 104, 20137–20141, doi:10.1073/pnas.0707383104 (2007).

37 Pobric, G., Jefferies, E. & Lambon Ralph, M. A. Amodal semantic representations depend on both anterior temporal lobes: Evidence from repetitive transcranial magnetic stimulation. Neuropsychologia 48, 1336–1342, doi:10.1016/j.neuropsychologia.2009.12.036 (2010).

38 Krizhevsky, A., Sutskever, I. & Hinton, G. E. in Advances in Neural Information Processing Systems 1097-1105 (Nevada, U.S.A, 2012).

39 He, K., Zhang, X., Ren, S. & Sun, J. Deep residual learning for image recognition. arXiv, doi: arXiv:1512.03385 (2015).

40 Chen, L., Lambon Ralph, M. A. & Rogers, T. T. A unified model of human semantic knowledge and its disorders. Nature Human Behaviour 1 (2017).

41 Kriegeskorte, N. Deep neural networks: A new framework for modeling biological vision and brain information processing. Annual Review of Vision Science 1, 417–446 (2015).

42 Kell, A. J. E., Yamins, D. L. K., Shook, E. N., Norman-Haignere, S. V. & McDermott, J. H. A task-optimized neural network replicates human auditory behaviour, predicts brain responses, and reveals a cortical processing hierarchy. Neuron 98, 630–644 (2018).

43 Plaut, D. C. Graded modality-specific specialisation in semantics: A computational account of optic aphasia. Cogn. Neuropsychol. 19, 603–639, doi:10.1080/02643290244000112 (2002).

44 Nelson, M. E. & Bower, J. M. Brain maps and parallel computers. Trends in Neurosciences 13, 403–408 (1990).

45 Kriegeskorte, N., Mur, M. & Bandettini, P. Representational Similarity Analysis - Connecting the branches of systems neuroscience. Frontiers in Systems Neuroscience 2 (2008).

46 Dilkina, K. & Lambon Ralph, M. A. Conceptual structure within and between modalities. Front. Hum. Neurosci. 31, 1–15, doi: doi: 10.3389/fnhum.2012.00333 (2013).

47 Cohen, J. D., Dunbar, K. & McClelland, J. L. On the control of automatic processes: A parallel distributed processing account of the Stroop effect. Psychol. Rev. 97, 332–361 (1990).

48 Morton, J. & Patterson, K. in Deep dyslexia (ed K. Patterson M. Coltheart, & J. C. Marshall) (Routledge & Kegan Paul, 1980).

49 Bozeat, S., Lambon Ralph, M. A., Patterson, K., Garrard, P. & Hodges, J. R. Non-verbal semantic impairment in semantic dementia. Neuropsychologia 38, 1207–1215 (2000).

50 Martin, A. GRAPES - Grounding representations in action, perception, and emotion systems: How object properties and categories are represented in the human brain. Psychon. Bull. Rev. 23, 979–990 (2016).

51 Bengio, Y. & Delalleau, O. in International Conference on Algorithmic Learning Theory. (eds Kivinen J., Szepesvári C., Ukkonen E., & Zeugmann T.) 18–36 (Springer).

52 Hochreiter, S. The Vanishing Gradient Problem During Learning Recurrent Neural Nets and Problem Solutions. International Journal of Uncertainty, Fuzziness and Knowledge-Based Systems 06, 107–116 (1998).

53 Saxe, A. M., McClelland, J. L. & Ganguli, S. Exact solutions to the nonlinear dynamics of learning in deep linear neural networks. arXiv, doi:arXiv:1312.6120 (2014).

54 Visser, M., Embleton, K. V., Jefferies, E., Parker, G. J. & Lambon Ralph, M. A. The inferior, anterior temporal lobes and semantic memory clarified: Novel evidence from distortion-corrected fMRI. Neuropsychologia, 48 (46) (pp 1689–1696), 2010, doi:http://dx.doi.org/10.1016/j.neuropsychologia.2010.02.016 (2010).

55 Rice, G. E., Hoffman, P. & Lambon Ralph, M. A. Graded specialization within and between the anterior temporal lobes. Annals of the New York Academy of Sciences 1359, 84–97 (2015).

56 Halai, A., Welbourne, S., Embleton, K. V. & Parkes, L. A comparison of dual-echo and spin-echo fMRI of the inferior temporal lobe. Human Brain Mapping 35, 4118–4128 (2014).

57 Chen, Y. et al. The ‘when’ and ‘where’ of semantic coding in the anterior temporal lobe: temporal representational similarity analysis of electrocorticogram data. Cortex 79, 1–13 (2016).

58 Marinkovic, K. et al. Spatiotemporal dynamics of modality-specific and supramodal word processing. Neuron 38, 487–497 (2003).

59 Herbet, G., Zemmoura, I. & Duffau, H. Functional anatomy of the inferior longitudinal fasciculus: From historical reports to current hypotheses. Frontiers in Neuroanatomy 12, doi:10.3389/fnana.2018.00077 (2018).

60 Bouhali, F. et al. Anatomical connections of the Visual Word Form Area. J. Neurosci. 34, 15402–15414 (2014).

61 Noonan, K. A., Jefferies, E., Visser, M. & Lambon Ralph, M. A. Going beyond inferior prefrontal involvement in semantic control: Evidence for the additional contribution of dorsal angular gyrus and posterior middle temporal cortex. J. Cogn. Neurosci. 25, 1824–1850 (2013).

62 McKee, J. L., Riesenhuber, M., Miller, E. K. & Freedman, D. J. Task dependence of visual and category representations in prefrontal and inferior temporal cortices. J. Neurosci. 34, 16065–16075 (2014).

63 Jackson, R. L., Cloutman, L. & Lambon Ralph, M. A. Exploring distinct default mode and semantic networks using a systematic ICA approach. Cortex 113, 279–297 (2019).

64 Davey, J. et al. Exploring the role of the posterior middle temporal gyrus in semantic cognition: Integration of anterior temporal lobe with executive processes. Neuroimage 137, 165–177 (2016).

65 Martin, A., Haxby, J. V., Lalonde, F. M., Wiggs, C. L. & Ungerleider, L. G. Discrete cortical regions associated with knowledge of color and knowledge of action. Science 270, 102–105, doi:10.1126/science.270.5233.102 (1995).

66 Chiou, R., Humphreys, G. F., Jung, J. & Lambon Ralph, M. A. Controlled semantic cognition relies upon dynamic and flexible interactions between the executive ‘semantic control’ and hub-and-spoke ‘semantic representation’ systems. Cortex 103, 100–116 (2018).

67 Binney, R. J., Parker, G. J. M. & Lambon Ralph, M. A. Convergent connectivity and graded specialization in the rostral human temporal lobe as revealed by diffusion-weighted imaging probabilistic tractography. J. Cogn. Neurosci. 24, 1998–2014 (2012).

68 Jung, J., Cloutman, L., Binney, R. J. & Lambon Ralph, M. A. The structural connectivity of higher order association cortices reflects human functional brain networks. Cortex 97, 221–239 (2016).

69 Rohde, D. L. T. LENS: The light, efficient network simulator. Technical Report CMU-CS-99-164, Carnegie Mellon University, Department of Computer Science, Pittsburgh, PA (1999).

70 IBM SPSS Statistics for Windows v. 25.0 (Armonk, NY, 2017).

71 Cloutman, L. L., Binney, R. J., Drakesmith, M., Parker, G. J. M. & Lambon Ralph, M. A. The variation of function across the human insula mirrors its pattern of structural connectivity: Evidence from in vivo probabilistic tractography. Neuroimage 59, 3514–3521 (2012).

72 McIntosh, A. R. Mapping cognition to the brain through neural interactions. Memory 7, 523–548, doi:10.1080/096582199387733 (1999).

