## Supplemental Materials for "Reverse-Engineering the Cortical Architecture for Controlled Semantic Cognition"

Supplementary Materials

Table of Contents

Supplementary Table 1	p1

Supplementary Table 2	p2

Supplementary Table 3	p3

Supplementary Table 4	p4

Supplementary Table 5	p4

Supplementary Table 6	p5

Supplementary Table 7	p5

Supplementary Note 1: Full Details of Contrasts and Size-Matching	p6

Supplementary Note 2:	Combining the Hidden Layers in a Single Similarity Metric	p7

Supplementary Note 3: Contrasting Similarity to Control and Representation	p8

Supplementary Note 4: Determining where Control Should Connect to the Model across all Architectures	p9

Supplementary Note 5: Modelling Distinct Representation & Control Impairments with Control Connected to the Spokes or Hidden Layer 1	p11

Supplementary Note 6: Assessing the Example Structure in each Region of Each Architecture in the Models with and without Control	p13

Supplementary Note 7: Similarity to the Multimodal and Unimodal Example Structures in the Models with and without Control	p15

Supplementary Note 8:	Varying the Sparsity of Shortcut Connections	p18

Supplementary Note 9: Similarity to the Context-Only and Context-Sensitive Example Structures	p20

Supplementary Methods 1: Matching the Number of Connections across Architectures	p22

Supplementary Methods 2: Simulation Code for LENS	p23

Supplementary Table 1. Summary of each metric per architecture and the effect of each contrast, in the models without control.

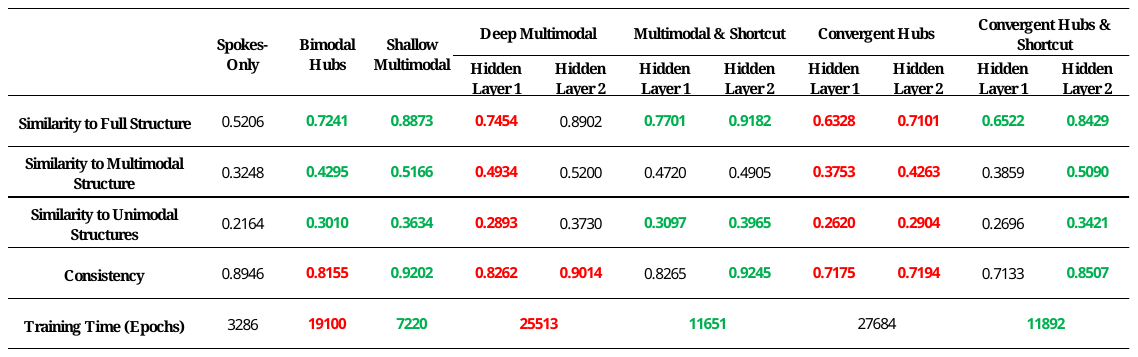

*Bold = significant change in the contrast of interest, resulting from the manipulation of a specific architectural feature. Green = significant improvement in the metric. Red = significant deterioration of the metric. The contrasts are the same as presented in text and compare each layer of each architecture to that layer in the prior architecture (where present), except the Convergent Hubs architecture, which is contrasted with the Deep Multimodal Hub architecture. Although also compared to the Multimodal Hub-plus-Shortcut architecture (see main text), the Convergent Hubs-plus-Shortcut architecture results are coloured based on the contrast with the Convergent Hubs architecture only. The greatest similarity to the full structure in a layer of an architecture represents that architectures conceptual abstraction score.*

Supplementary Table 2. Summary of each metric per architecture and the effect of each contrast, in the models with control.

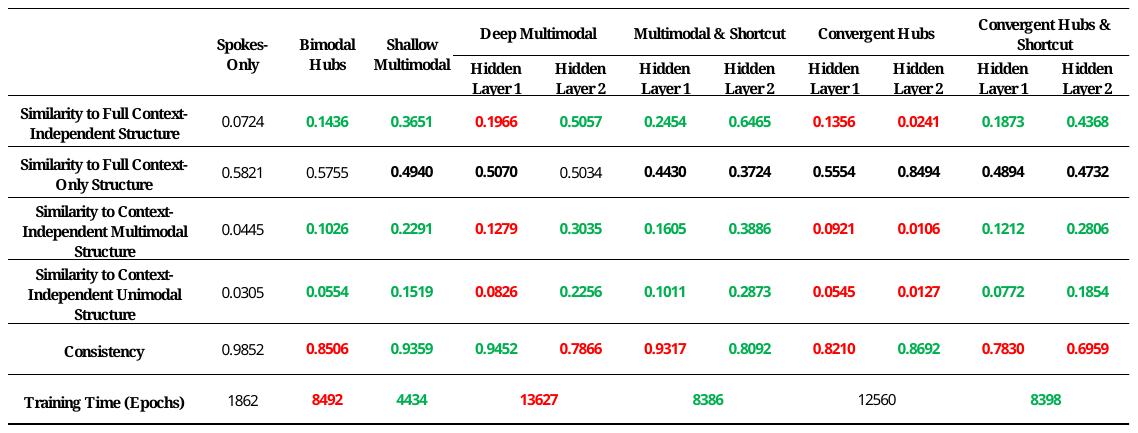
 *Bold = significant change in the contrast of interest, resulting from the manipulation of a specific architectural feature. Green = significant improvement in the metric. Red = significant deterioration of the metric. As the similarity to the full control structure is not intrinsically positive or negative, instead depending on the effect on the similarity to the representation structure, significant changes in this metric are shown in bold but are not coloured. The contrasts are the same as presented in text and compare each layer of each architecture to that layer in the prior architecture (where present), except the Convergent Hubs architecture, which is contrasted with the Deep Multimodal Hub architecture. Although also compared to the Multimodal Hub-plus-Shortcut architecture (see main text), the Convergent Hubs-plus-Shortcut architecture results are coloured based on the contrast with the Convergent Hubs architecture only. The greatest similarity to the full structure in a layer of an architecture represents that architectures conceptual abstraction score.*

Supplementary Table 3. The beta weights and significance of each predictor (the similarity matrix from each area) in the regression model for each architecture when predicting the Context-Independent example structure.

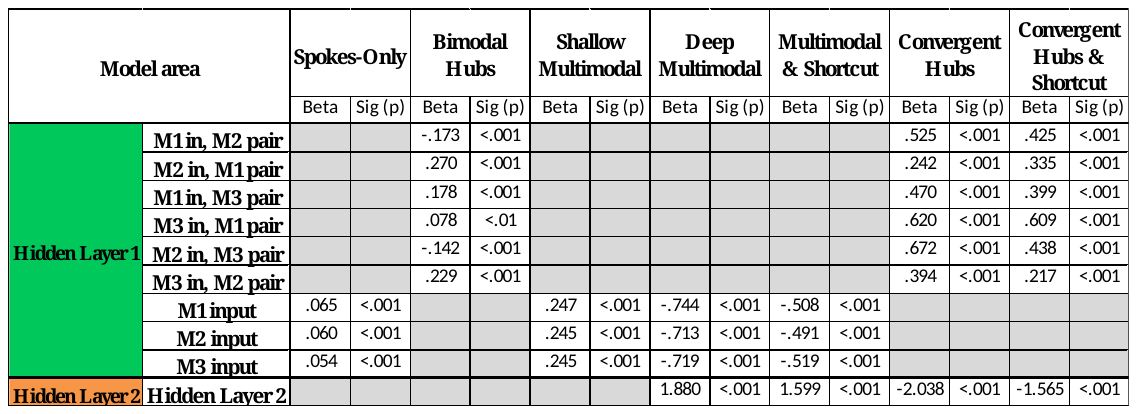

*M1 = modality 1, M2 = modality 2, M3 = modality 3. Sig = significance.*

Supplementary Table 4. The single closest orthogonal example structure to the representations in each area of each architecture without control.

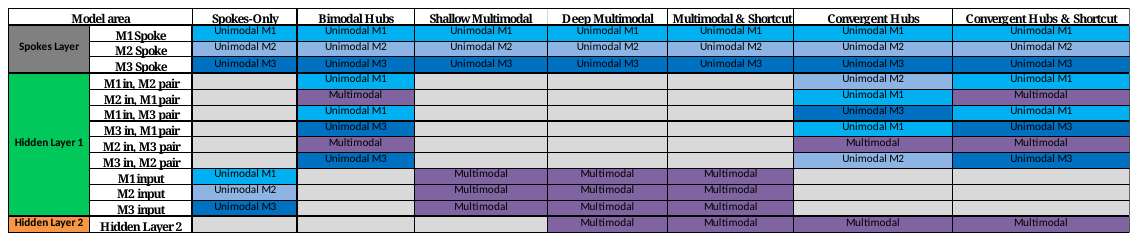
*M1 = modality 1, M2 = modality 2, M3 = modality 3.*

Supplementary Table 5. The closest combination of the orthogonal example structures to the representations in each area of each architecture without control.

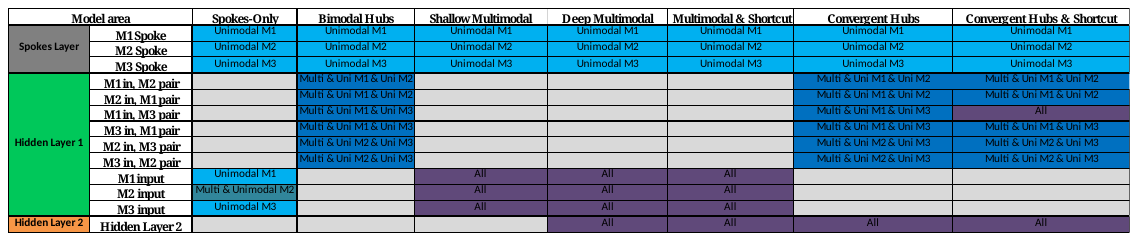

*M1 = modality 1, M2 = modality 2, M3 = modality 3. Uni = unimodal, Multi = multimodal. All = multimodal, unimodal M1, unimodal M2 & unimodal M3.*

Supplementary Table 6. The single closest orthogonal Context-Independent example structure to the representations in each area of each architecture with control.

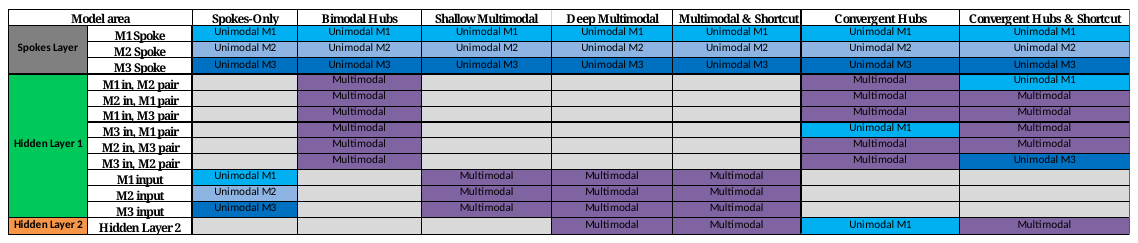

*M1 = modality 1, M2 = modality 2, M3 = modality 3.*

Supplementary Table 7. The closest combination of the orthogonal Context-Independent example structures to the representations in each area of each architecture with control.

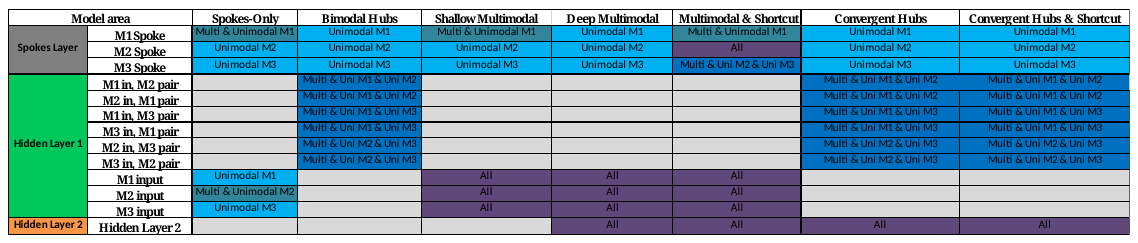

*M1 = modality 1, M2 = modality 2, M3 = modality 3. Uni = unimodal, Multi = multimodal. All = multimodal, unimodal M1, unimodal M2 & unimodal M3.*

Supplementary Note 1: Full Details of Contrasts and Size-Matching

As the main text focuses on the conceptual abstraction score derived from the highest layer of each architecture, contrasts of the similarity to the Context-Independent structure in the layer with less conceptual abstraction (typically Hidden Layer 1) are not displayed for the deeper models. The architectures in Phase 1 (where Hidden Layer 2 always showed the greatest abstraction), varied significantly on conceptual abstraction within Hidden Layer 1 (F(6, 2393)=826.841, p<.001). As well as the importance of a multimodal hub reported in text, negative effects were found for depth (*Shallow Multimodal Hub > Deep Multimodal Hub;* t(478)=36.357, p<.001) and hierarchical convergence (*Deep Multimodal Hub > Convergent Hubs*; t(718)=22.378, p<.001; *Multimodal Hub-plus-Shortcut* *> Convergent Hubs-plus-Shortcut*; t(718)=23.889, p<.001). Shortcut connections resulted in greater conceptual abstraction in Hidden Layer 1 (*Deep Multimodal Hub vs. Multimodal Hub -plus-Shortcut*; t(478)=-4.833, p<.001; *Convergent Hubs* *vs. Convergent Hubs-plus-Shortcut*; t(958)=-4.008, p<.001).

In Phase 2, conceptual abstraction was greatest in Hidden Layer 2 for all architectures except the *Convergent Hubs* architecture. The architectures varied on the conceptual abstraction of information represented in Hidden Layer 1 (F(6, 2393)=780.659, p<.001, Figure 3). As in Phase 1, a multimodal hub and shortcut connections *(Deep Multimodal Hub < Multimodal Hub-plus-Shortcut*; t(454.479)=-15.637, p<.001; *Convergent Hubs* *< Convergent Hubs-plus-Shortcut*; t(852.82)=-11.331, p<.001) led to greater Context-Independent structure in Hidden Layer 1, whereas this structure was reduced by depth (*Shallow Multimodal Hub* *> Deep Multimodal Hub*; t(430.156)=50.150, p<.001) and hierarchical convergence (*Deep Multimodal Hub* *>* *Convergent Hubs*; t(715.896)=18.809, p<.001; *Multimodal Hub-plus-Shortcut > Convergent Hubs-plus-Shortcut*; t(714.279)=12.994, p<.001). Within Hidden Layer 2, hierarchical convergence also reduced conceptual abstraction (*Deep Multimodal Hub >* *Convergent Hubs*; t(99.952)=46.962) although this was improved by the addition of shortcut connections (*Convergent Hubs* *< Convergent Hubs-plus-Shortcut*; t(94.394)=-34.918).

For the across-architecture contrasts of similarity to the examples structures, the set of hidden units included are defined by their coherence, i.e., the units which all have the same potential connections. In some cases for the contrasts of the similarity in Hidden Layer 1, these regions contain different numbers of units, which could affect this similarity. Therefore, where possible additional size-matched contrasts were employed to check the results of the main contrast. Specifically, in architectures *Bimodal Hubs*, *Convergent Hubs*, *Convergent Hubs-plus-Shortcut,* the full bimodal hubs were used. This allowed size-matched contrasts between *Spokes-Only* and *Bimodal Hubs*, *Bimodal Hubs* and *Shallow Multimodal Hub*, *Deep Multimodal Hub* and *Convergent Hubs* and *Multimodal Hub-plus-Shortcut* and *Convergent Hubs-plus-Shortcut*. In Part I, all results were confirmed with size-matched versions (*Spokes-Only vs. Bimodal Hubs*; t(435.331)=-54.317, p<.001; *Bimodal Hubs vs. Shallow Multimodal Hub*; t(375.426)=-23.774, p<.001; *Multimodal Hub-plus-Shortcut vs. Convergent Hubs-plus-Shortcut*; t(478)=3.837, p<.001) except for the significant decrease with hierarchical convergence in Hidden Layer 1 (*Deep Multimodal Hub vs. Convergent Hubs*; t(474.563)=-1.404, p=.161). Although this effect may partially relate to size, the effect of hierarchical convergence is still significant when comparing the architectures with shortcut connections after matching for size. In Part II, size-matched contrasts supported all findings reported in the main text, including the detrimental effect of hierarchical convergence (*Spokes-Only vs. Bimodal Hubs*; t(284.424)=-27.278, p<.001; *Bimodal Hubs vs. Shallow Multimodal Hub*; t(478)=-52.294, p<.001; *Deep Multimodal Hub vs.* *Convergent Hubs*; t(397.594)=11.729, p<.001; *Multimodal Hub-plus-Shortcut vs. Convergent Hubs-plus-Shortcut*; t(347.873)=5.206, p<.001). No size-matched contrast is possible between the *Shallow Multimodal Hub* and *Deep Multimodal Hub* architectures, or for the contrasts between Hidden Layer 1 in the *Convergent Hubs* architecture and Hidden layer 2 in the *Deep Multimodal Hub* and *Convergent Hubs-plus-Shortcut* architectures due to the different total size of Hidden Layer 1.

Supplementary Note 2: Combining the Hidden Layers in a Single Similarity Metric

The conceptual abstraction score is computed using the similarity to the example structure in each set of coherent units in the deepest layer (i.e., Hidden Layer 2 or the average across 3 or 6 coherent regions of Hidden Layer 1, see Methods). Thus, it is assumed that the architecture that has the region closest to an example structure is equivalent to the architecture best able to represent the structure overall. However, it may be that some architectures represent more or less overlapping structure in different regions and therefore the total structure represented is greater in an architecture with lower similarity per region. This assumption may be tested by creating a regression model per architecture that takes into account the similarity matrix from all parts of the model (i.e., 3 or 6 Hidden Layer 1 regions, plus Hidden Layer 2 where present) and flexibly weights them in order to best predict the example structure. Thus, for each pair of examples the regression model predicts their underlying conceptual similarity using the similarity of the activation values in each section of Hidden Layer 1 and 2. The amount of variance explained by each architecture overall (R²) can then be used as a metric of how well the architecture represents the example structure across all hidden units and allows (non-statistical) comparison of the different architectures.

The structure in the constituent regions of each architecture was significantly predictive of the context-independent semantic representations *(*Supplementary Figure 1; *Spokes-Only;* F(3, 10292)=45.021, p<.001; *Bimodal Hubs;* F(6, 10289)=91.120, p<.001; *Shallow Multimodal Hub;* F(3, 10292)=1363.052, p<.001; *Deep Multimodal Hub;* F(4, 10291)=7733.241, p<.001; *Convergent Hubs;* F(7, 10288)=993.382, p<.001; *Multimodal Hub-plus-Shortcut;* F(4, 10291)=18486.250, p<.001; *Convergent Hubs-plus-Shortcut;* F(7, 10288)=1436.551, p<.001). All regions contributed to each architectures prediction although not to an equal extent (in particular, wherever present the deep hub had the greatest absolute weight; Supplementary Table 3). However, dramatic differences were found in the models ability to extract the Context-Independent structure, with the activity in the ‘best’ architecture (*Multimodal Hub-plus-Shortcut* ) achieving 87.7% variance explained, and the ‘worst’ architecture (*Spokes-Only)* only able to explain 1.3%. Greater variance was explained with a hub (*Spokes-Only* R²= .013, *Bimodal Hubs* R²= .050), with a multimodal hub (*Bimodal Hubs* R²= .050, *Shallow Multimodal Hub* R²= .284), with depth (*Shallow Multimodal Hub* R²= .284, *Deep Multimodal Hub* R²= .750) and with shortcut connections (*Deep Multimodal Hub* R²= .750, *Multimodal Hub-plus-Shortcut* R²= .878; *Convergent Hubs* R²= .403, *Convergent Hubs-plus-Shortcut* R²= .494). Less variance was explained when the architecture included hierarchical convergence (*Deep Multimodal Hub* R²= .750, *Convergent Hubs* R²= .403; *Multimodal Hub-plus-Shortcut* R²= .878*, Convergent Hubs-plus-Shortcut* R²= .494). Therefore, using the regression method shows the same across-architecture pattern as assessing the similarity of the representations in each area separately, confirming the viability of the current assessment method. Although not shown, the across-architecture pattern found using regression analyses for the multimodal and unimodal Context-Independent structures and the full Context-Only structure also mirrored the architectures with the single highest most similar area.

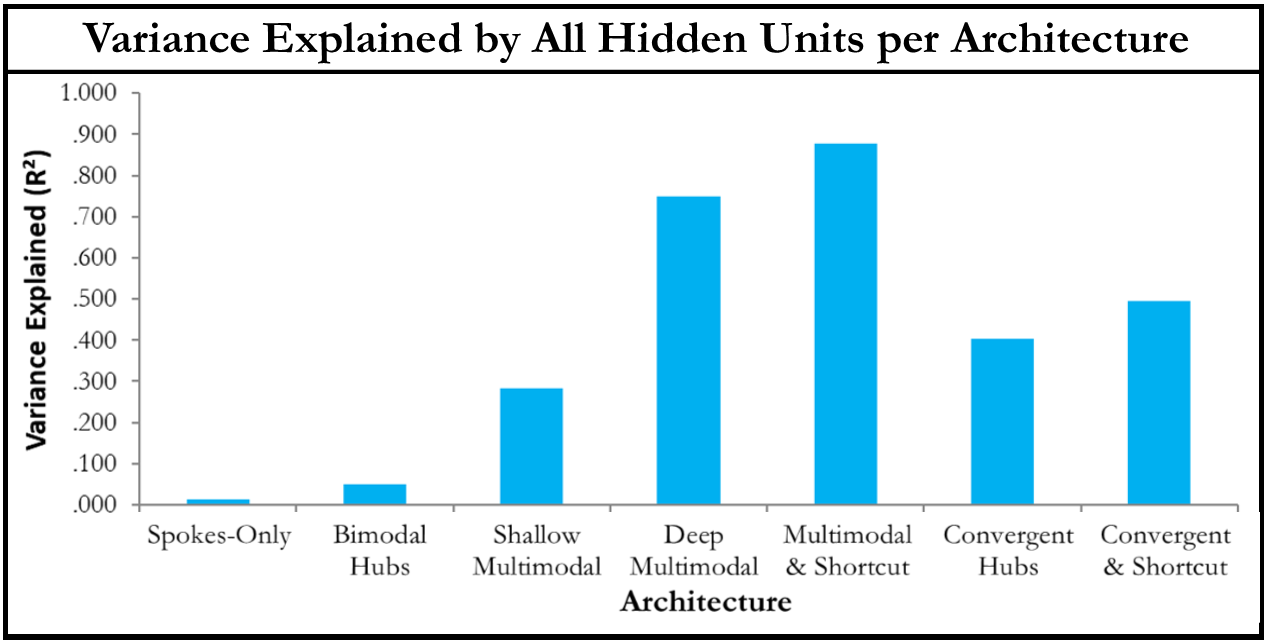

*Supplementary Figure 1. How well the representations in each architecture could explain the full Context-Independent examples structure in a regression analysis. For each architecture the variance in the examples structure that can be explained using any combination of the sets of hidden units is shown as R². This analysis incorporates all hidden layers within a model and therefore compares models fairly regardless of which layer is a better estimate of the examples structure.*

Supplementary Note 3: Contrasting Similarity to Control and Representation

In the Phase 2 simulations employing context-sensitivity, any given region of an architecture could be of greater importance for control or representation. To assess this relative importance the abstraction of the context-independent semantic representation structure may be directly compared to the extraction of the control signal via paired t-tests. Only the deep hub in the *Multimodal Hub-plus-Shortcut* architectures showed a significant preference for the Context-Independent structure (Context-Independent *>* Control Only; Hidden Layer 2; *Multimodal Hub-plus-Shortcut;* t(158)=23.883, p<.001). The hub in the *Deep Multimodal Hub* architecture has no significant preference (Context-Independent *~=* Control Only; Hidden Layer 2; *Deep Multimodal Hub;* t(158)=0.164, p=.870). All other regions and architecture expressed a clear preference for the control structure (Context-Independent *<* Control Only; Hidden Layer 1; *Spokes-Only*; t(415.821)=-312.491, p<.001; *Bimodal Hubs;* t(932.181)=-122.86, p<.001; *Shallow Multimodal Hub;* t(461.228)=-36.244, p<.001; *Deep Multimodal Hub;* t(478)=-107.049, p<.001; *Convergent Hubs;* t(903.273)=-99.291, p<.001; *Multimodal Hub-plus-Shortcut;* t(478)=-55.902, p<.001; *Convergent Hubs-plus-Shortcut;* t(958)=-54.101, p<.001; Hidden Layer 2; *Convergent Hubs;* t(121.836)=-111.311, p<.001; *Convergent Hubs-plus-Shortcut;* t(158)=-2.215, p<.001). Thus, a relative functional specialisation for representation processes begins to emerge in the deep hub only when there is a single, deep multimodal hub and is supported by the presence of shortcut connections. This functional specialisation may be critical for the conceptual abstraction performance criterion, optimal in the *Multimodal Hub-plus-Shortcut* architecture.

Performance of the same analysis on the simulations with selective connections from control to each layer, allow assessment of the effect of these connections on the relative functional specialisation of the hub. This relative specialisation is maintained with connections to the Spokes Layer (Context-Independent > Context-Only; t(158)=76.211, p<.001) or Hidden Layer 1 (Context-Independent > Context-Only; t(142.748)=45.454, p<.001). However, if control is connected to the deep hub only, this specialisation is not possible, and the hub is instead specialised for control (Context-Independent < Context-Only; t(158)=-21.964, p<.001). In all cases Hidden Layer 1 is closest to the Context-Only structure (Context-Independent vs Context-Only; Spokes Layer control connection; t(466.244)=-44.903, p<.001; Hidden Layer 1 control connection; t(478)=-86.395, p<.001; Hidden Layer 2 control connection; t(431.159)=-82.908, p<.001). Thus, the resulting reduction in conceptual abstraction when control is connected directly to the deep hub co-occurs with the lack of functional specialisation of this region for representation.

Supplementary Note 4: Determining where Control Should Connect to the Model across all Architectures

Assessing the absolute values of the emergent weights from control to each layer demonstrated similar results across all deep architectures whereby there was greater reliance on the peripheral layers than the deep hub (Supplementary Figure 2; Spokes Layer *>* Hidden Layer 2; *Deep Multimodal Hub;* t(1358.735)=9.944, p<.001; *Convergent Hubs;* t(1447.373)=11.897, p<.001; *Convergent Hubs-plus-Shortcut;* t(1486.656)=11.240, p<.001; Hidden Layer 1 *>* Hidden Layer 2; *Deep Multimodal Hub;* t(1503.101)=12.029, p<.001; *Convergent Hubs;* t(1589.941)=20.280, p<.001; *Convergent Hubs-plus-Shortcut;* t(1549.674)=16.297, p<.001). Less consistency was apparent in the reliance on the connections to Spokes Layer compared to Hidden Layer 1, with architectures with bimodal hubs relying on Hidden Layer 1 more (Spokes Layer *<* Hidden Layer 1; *Bimodal Hubs;* t(3028.887)=-6.869, p<.001; *Convergent Hubs;* t(2115.817)=-5.193, p<.001; *Convergent Hubs-plus-Shortcut;* t(2129.273)=-3.152, p<.005) and the *Spokes-Only* architecture showing the opposite pattern (Spokes Layer *>* Hidden Layer 1; t(2971.586)=5.474, p<.001). No other architecture displays a preference (Spokes Layer *~=* Hidden Layer 1; *Shallow Multimodal Hub;* t(3118.939)=1.413, p=.158; *Deep Multimodal Hub;* t(2127.407)=-1.304, p=.577).

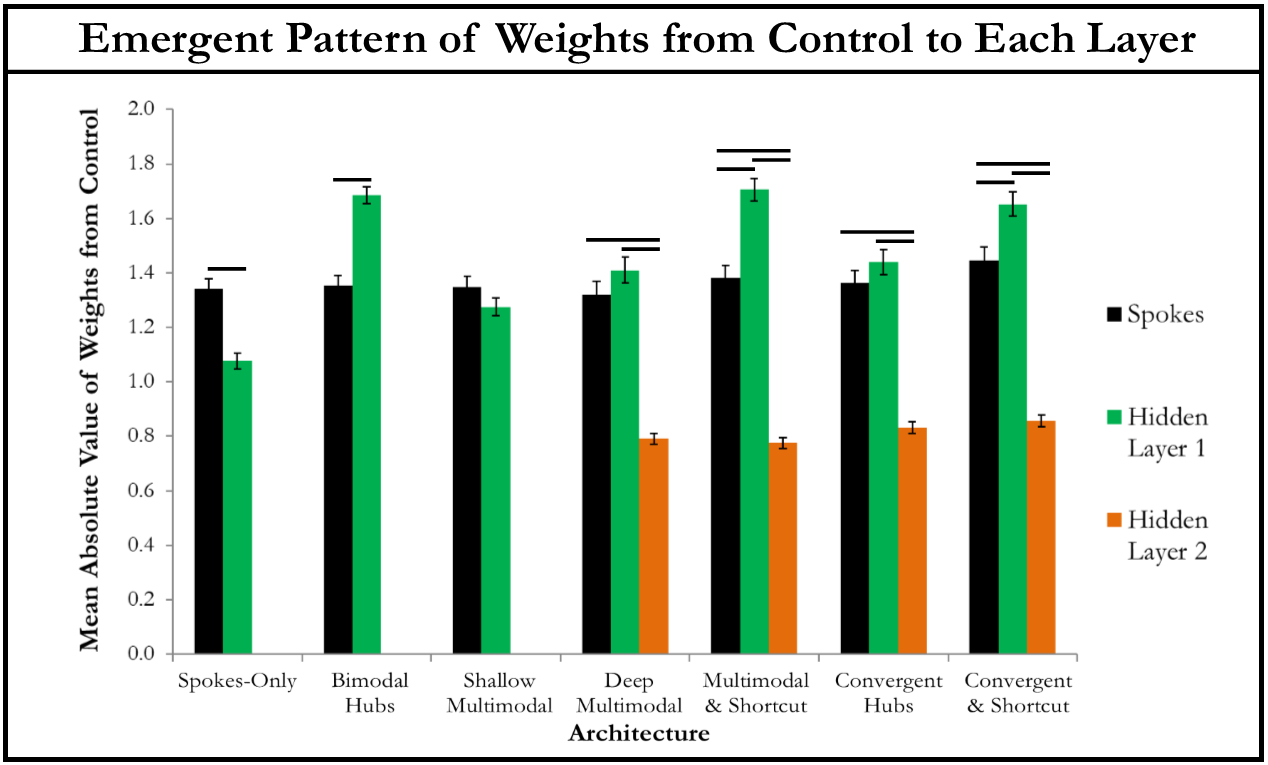
*Supplementary Figure 2. Comparing the absolute value of the weights from control to each layer of the model across all architectures. Significant differences between layers are highlighted with a black line.*

All deep architectures had greater conceptual abstraction scores when control was connected to peripheral regions than the deep hub (Supplementary Figure 3.A.; Spokes Layer *>* Hidden Layer 2; *Deep Multimodal Hub;* t(125.667)=46.182, p<.001; *Convergent Hubs;* t(85.725)=, p<.001; *Convergent Hubs-plus-Shortcut;* t(158)=44.172, p<.001; Hidden Layer 1 *>* Hidden Layer 2; *Deep Multimodal Hub;* t(158)=32.050, p<.001; *Convergent Hubs;* t(87.688)=19.247, p<.001; *Convergent Hubs-plus-Shortcut;* t(137.896)=33.458, p<.001). Again, the difference between connections to the Spokes Layer and Hidden Layer 1 was mixed (Spokes Layer *>* Hidden Layer 1; *Deep Multimodal Hub;* t(127.324)=7.347, p<.001; *Convergent Hubs;* t(158)=22.878, p<.001; *Convergent Hubs-plus-Shortcut;* t(158)=3.816, p<.001; Spokes Layer *<* Hidden Layer 1; *Spokes-Only;* t(478)=-12.362, p<.001; *Bimodal Hubs;* t(958)=10.449, p<.001; Spokes Layer *>* Hidden Layer 1; Spokes Layer *~=* Hidden Layer 1; *Shallow Multimodal Hub;* t(478)=1.452, p=.147).

Conceptual abstraction was also greater in Hidden Layer 1 if control was connected to the Spokes than Hidden Layer 2 (Spokes Layer *>* Hidden Layer 2; *Deep Multimodal Hub;* t(478)=8.335, p<.001; *Convergent Hubs;* t(940.258)=14.450, p<.001; *Convergent Hubs-plus-Shortcut;* t(958)=6.578, p<.001), although a connection to Hidden Layer 1 was not always preferable (Hidden Layer 1 *>* Hidden Layer 2; *Deep Multimodal Hub;* t(478)=6.164, p<.001; Hidden Layer 1 *~=* Hidden Layer 2; *Convergent Hubs;* t(958)=1.410, p=.476; *Convergent Hubs-plus-Shortcut;* t(958)=1.366, p=.517). The connection to the Spokes Layer was preferred for architectures with bimodal hubs only (Spokes Layer *~=* Hidden Layer 1; *Deep Multimodal Hub;* t(478)=1.715, p=.261; Spokes Layer *>* Hidden Layer 1; *Convergent Hubs;* t(942.686)=13.076, p<.001; *Convergent Hubs-plus-Shortcut;* t(958)=5.019, p<.001). The time taken to train each version of each architecture is shown in Supplementary Figure 3.D. Differences in this measure were subtle, with no difference reaching p<.001. Regardless of the specific architecture, conceptual abstraction was greater with control connected to peripheral regions and not a deep hub.

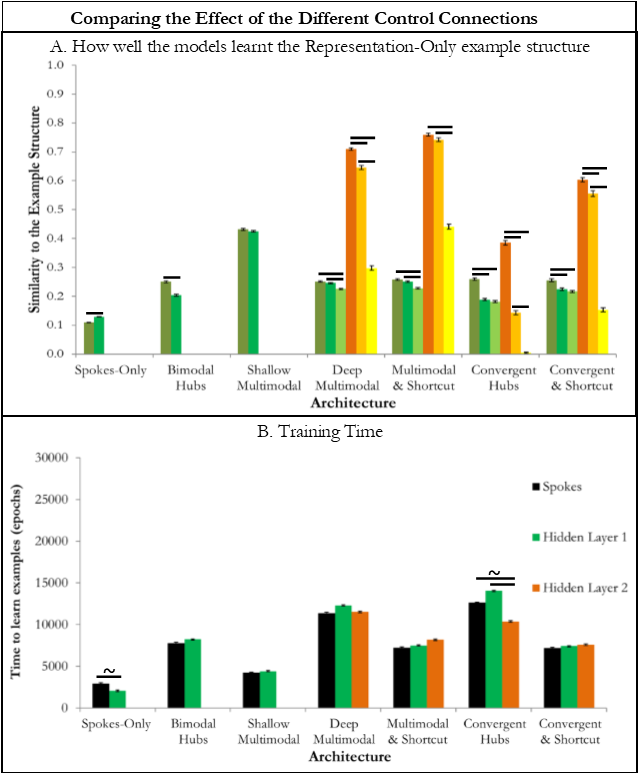

*Supplementary Figure 3. Comparing the effect of connecting the control units to each of the three different layers in each architecture. A. The similarity between the Context-Independent example structure and the representations in Hidden Layer 1 (green variants) and Hidden Layer 2 (orange variants) with the control signal connected to the Spokes Layer (darkest green/orange), Hidden Layer 1 (mid green/orange) or Hidden Layer 2 (lightest green/orange). The bar shows the mean similarity value across the different runs of the model (bars show standard error). Significant differences in the planned contrasts are highlighted with a black line (p<.05). B. The time taken to learn the examples is shown as the number of epochs of training with control connected to the Spokes Layer (black), Hidden Layer 1 (green) or Hidden Layer 2 (orange). The bars show the mean number of epochs across different runs of the model (error bars show standard error). Significant differences in the planned contrasts are highlighted with a line (p<.05). A trend (p<.1) is highlighted with a tilde.*

Supplementary Note 5: Modelling Distinct Representation & Control Impairments with Control Connected to the Spokes or Hidden Layer 1

Lesion analyses were performed in the main text for the version of the *Multimodal Hub-plus-Shortcut* architecture with the Control Layer connected to the Spokes Layer. For the errors that were found to significantly vary by damage type (i.e., inappropriate and appropriate commissions) planned contrasts were used to compare the number of errors at each damage level (except no damage) between the Control and Representation Damage types. Significant differences were found between the damage types at every level for both incorrect (Level 1; t(38)=-15.849, p<.001; Level 2; t(29.285)=-47.766, p<.001; Level 3; t(38)=-25.071, p<.001; Level 4; t(38)=-22.284, p<.001) and correct feature-type commissions (Level 1; t(19.687)=13.095, p<.001; Level 2; t(19.413)=17.024, p<.001; Level 3; t(19.397)=20.357, p<.001; Level 4; t(19.981)=20.954, p<.001).

In addition, lesion analyses may be performed for the version of the *Multimodal Hub-plus-Shortcut* architecture with the Control Layer connected to Hidden Layer 1. Levels of damage with equivalent numbers of errors were identified. This resulted in Representation Damage simulations with the removal of connections at proportions of 0, 0.1, 0.15, 0.2 and 0.35. The Control Damage versions had noise added to the control signal at a range of 0, 0.49375, 0.625, 0.75 and 1. The number of errors of each type is shown for each damage type in Supplementary Figure 4, with the total height of the bar dictating the total number of errors.

Unlike the simulations where control was connected to the spoke regions, there was a significant effect of damage type, as well as damage level, on the number of omission errors (damage type; F(1,190)=17.770, p<.001; level; F(4,190)=17.770, p<.001; interaction; F(4,190)=10.065, p<.001). However, very few direct comparisons between the damage types reached significance with only the final two levels significantly higher for Control Damage than Representation Damage (Representation Damage vs. Control Damage; Level 1; t(38)=0.868, p=.391; Level 2; t(38)=-0.017, p=.986; Level 3; t(31.731)=-2.561, p=.015; Level 4; t(38)=-5.626, p<.001). These differences are relatively minor. Greater differences were identified for Correct Feature-Type Commissions and Incorrect Feature-Type Commissions, as in the simulations with control connected to the Spokes Layer. Incorrect Feature-Type Commissions are significantly affected by damage type, damage level and their interaction (damage type; F(1,190)=2496.052, p<.001; damage level; F(4, 190)=247.700, p<.001; interaction; F(4, 190)=181.188, p<.001). These context-inappropriate commissions are significantly higher following damage to control (Representation Damage vs. Control Damage; Level 1; t(38)=-25.162, p<.001; Level 2; t(22.976)=-26.898, p<.001; Level 3; t(32.230)=-25.211, p<.001; Level 4; t(38)=-24.811, p<.001). Damage type and damage level are also significant for the Correct Feature-Type Commissions (damage type; F(1, 190)=1095.371, p<.001; level; F(4,190)=113.262, p<.001; interaction; F(4,190)=10.065, p<.001). These commissions are significantly higher following representation damage (Representation-Damage vs. Control-Signal-Damage; Level 1; t(19.686)=14.552, p<.001; Level 2; t(20.344)=19.465, p<.001; Level 3; t(19.341)=13.925, p<.001; Level 4; t(19.476)=20.325, p<.001).

Thus, whether control is connected to the Spokes Layer or Hidden Layer 1, the Representation-Damage led to a greater number of Correct Feature-Type Commissions, whereas both types of control damage led to an increase in Incorrect Feature-Type Commissions. Although qualitatively similar, the Control-Signal-Damage version led to a greater increase in Incorrect Feature-Type Commissions. Therefore, damage to regions with greater responsibility for representation and control led to qualitatively different patterns of impairment.

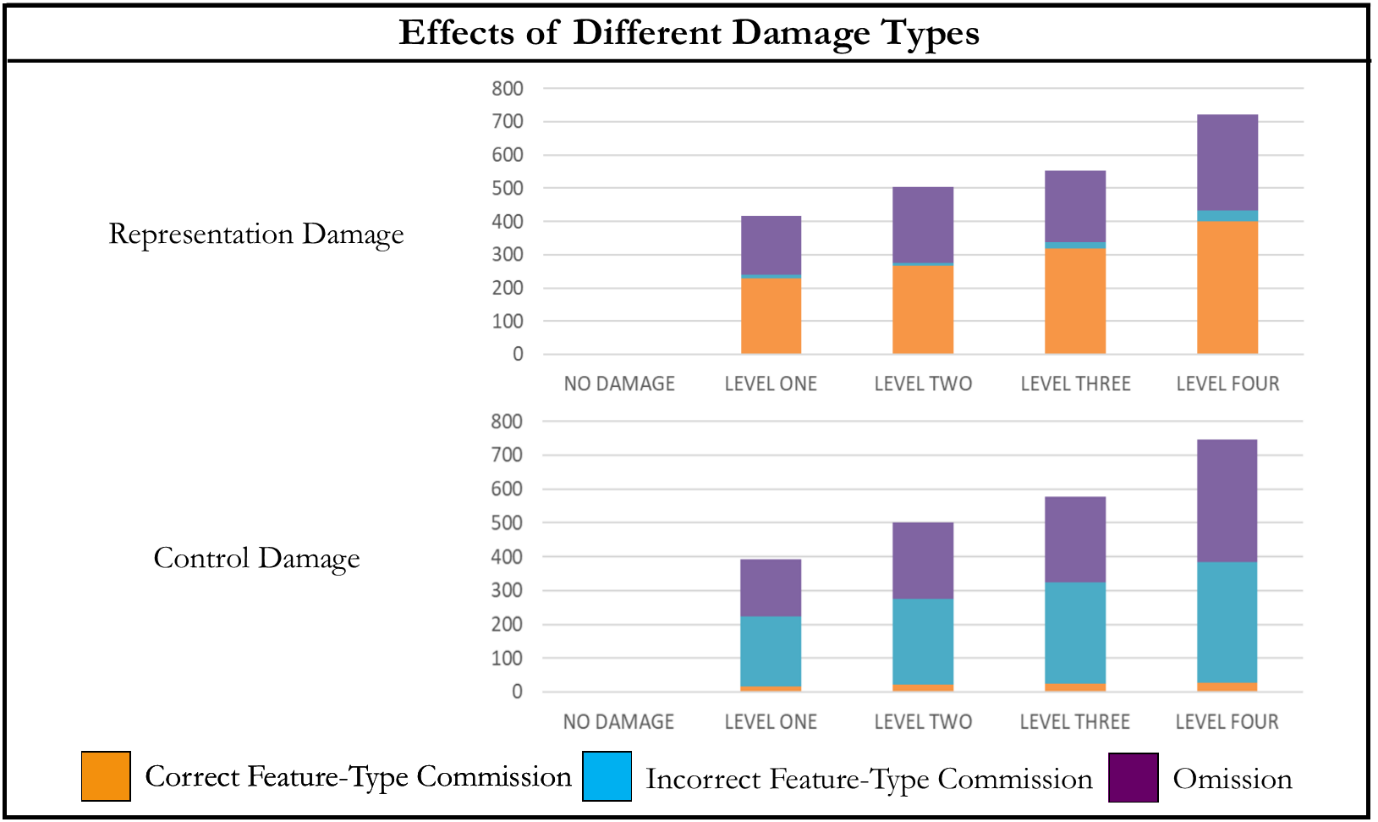

Omission

*Supplementary Figure 4. The effects of damage to control and representation within the Multimodal Hub-plus-Shortcut architecture when control is connected to Hidden Layer 1. The different kinds of feature errors resulting from the three types of damage are shown. The height of the bar shows the number of errorful features, which is matched across damage types at each level. Each errorful feature is an omission of a correct features (purple), a commission of a feature of the correct type (orange), or a commission of a feature of the incorrect type (blue).*

Supplementary Note 6: Assessing the Example Structure in each Region of Each Architecture in the Models with and without Control

The full example structure was constructed using four orthogonal structures; multimodal, unimodal M1, unimodal M2 and unimodal M3. In order to consider how the different regions and architectures process these multimodal and unimodal example structures, the example structure closest to the representations in each area was determined based on the correlation between the similarity matrices. The areas assessed include each modality-specific region in the Spokes Layer (labelled by modality; M1 Spoke, M2 Spoke and M3 Spoke), each coherent area within Hidden Layer 1, and Hidden Layer 2. In an architecture including bimodal hubs, Hidden Layer 1 consists of 6 different regions, each including input from one modality-specific spoke, and connections to an equivalent region with a different modality of input. These regions are labelled based on the modality of input and the modality of input of their paired region (e.g., M2 input, paired with M3). In an architecture without bimodal hubs, Hidden Layer 1 consists of 3 coherent regions, one per modality of input (e.g., M2 input). The similarity matrix across examples was constructed for each distinct region of each architecture and compared to each of the orthogonal example structures (multimodal, unimodal M1, unimodal M2 and unimodal M3). The numerically closest single example structure is displayed for each region of the simulations without control in Supplementary Table 4. Then, the similarity between the representation in each region and every possible combination of these orthogonal structures was assessed. The closest combination of example structures per region is displayed in Supplementary Table 5. The colours of the cells help show the pattern of representations across regions and architectures, reflecting the four structures in Supplementary Table 4 and the number of combined structures in Supplementary Table 5 (i.e., a single unimodal structure is light blue, whereas the full combination of the 3 unimodal and the multimodal structure is purple and intermediate combinations are represented with darker blues).

Overall, the spoke regions were always closest to the single unimodal structure present in their respective modalities. The 3 sections of Hidden Layer 1 in the *Spokes-Only* architecture also represented each of the 3 unimodal structures, although one section was closer to the combination of its respective unimodal structure with the multimodal structure. Regardless of specific architecture, the representations in the bimodal hubs were typically closest to a unimodal structure. However, when the combination of structures was assessed these hubs were shown to combine the two relevant unimodal structures and the multimodal structure i.e., to represent the full information available across two modalities of input. Thus, these regions successfully act as bimodal hubs integrating pairs of modalities. With only bimodal hubs there is no region that is closest to representing the full combination of all the orthogonal structures. The single closest structure in the multimodal hubs is the multimodal structure, which is the most informative as to the full similarity structure of the concepts. Overall, these multimodal hubs represent the full combination of structures (i.e. the multimodal and all 3 unimodal structures). This is true whether the multimodal hub is in Hidden Layer 1 or 2, i.e., in both the shallow and deep architectures. The deep architectures that do not have bimodal hubs (i.e., the *Deep Multimodal Hub* and *Multimodal Hub-plus-Shortcut* architectures) have distinct routes to the multimodal hub for information in each modality. However, interestingly these Hidden Layer 1 regions are also found to best represent the multimodal structure and to reflect the combination of all the orthogonal structures. Note that without the multimodal hub the connections to this layer from the Spokes Layer would be qualitatively equivalent to the *Spokes-Only* architecture. Thus, the presence of the deep multimodal hub changes the nature of the representations at shallower levels.

The same investigation was performed for the simulations with control. The orthogonal structures are computed using the Context-Independent structure and are therefore independent of context. The closest single structure is displayed in Supplementary Table 6 and the closest combination in Supplementary Table 7. The single closest structure is similar to without control for all regions except the bimodal hubs which are typically closest to the multimodal structure. However, these hubs are still closest to the combination of two unimodal structures plus the multimodal structure, i.e., the information that it is possible to gain from input in two modalities. Hidden Layer 2 of the *Convergent Hubs* architecture is closest to a unimodal structure, although all deep hubs still reflect the full combined structure. The results of assessing the closest combined structure in the Spokes Layer is more diverse than without control. In some cases the Spokes Layer is closer to the combination of a unimodal and the multimodal structure. In the *Multimodal Hub-plus-Shortcut* architecture, the spokes are closest to different combinations of example structures. This may suggest a change in the representations stored in the Spokes Layer when shortcut connections are added to a model with a controlled output.

Supplementary Note 7: Similarity to the Multimodal and Unimodal Example Structures in the Models with and without Control

Assessing the ability of the models to extract the full example structure is the primary aim of this work. However, this structure consists of orthogonal multimodal and unimodal structures. For a simulation to acquire either requires conceptual abstraction across time, yet only the (more informative) multimodal structure necessitates abstraction across modalities. Thus, to assess the relative abilities of the different architectures to integrate information across the different modalities it may be useful to specifically assess how well an architecture can represent the multimodal structure (*vs.* the unimodal structure). In the simulations without control, conceptual abstraction of the multimodal structure varied by architecture (Supplementary Figure 5.A; Hidden Layer 1; F(6,2393)=69.167, p<.001; Hidden Layer 2; F(3,316)=9.941, p<.001, all significant contrasts had p<.001 unless stated) and was improved by the addition of a hub (*Spokes-Only < Bimodal Hubs*; t(667.329)=-12.060), a multimodal hub (*Bimodal Hubs < Shallow Multimodal Hub*; t(709.582)=-10.738, p<.001) and shortcut connections in the presence of bimodal hubs only (*Deep Multimodal Hub ~= Multimodal Hub-plus-Shortcut*; t(158)=2.062, p=.163; *Convergent Hubs< Convergent Hubs-plus-Shortcut*; t(158)=-3.571, p<.005). Hierarchical convergence significantly decreased conceptual abstraction only when shortcut connections were present (*Deep Multimodal Hub > Convergent Hubs*; t(125.334)=5.058; *Multimodal Hub-plus-Shortcut ~= Convergent Hubs-plus-Shortcut*; t(136.68)=-0.926, p=1). Depth had no significant effect (*Shallow Multimodal Hub ~= Deep Multimodal Hub*; t(318)=-0.339, p=.735). Thus, the multimodal structure displayed a very similar profile to the full structure. Indeed, the unimodal structure was similarly affected by architecture (Supplementary Figure 5.B.; Hidden Layer 1; F(6,7193)=41.974, p<.001; Hidden Layer 2; F(3,956)=33.990, p<.001) with positive effects of a hub (*Spokes-Only < Bimodal Hubs*; t(1156.15)=-7.118), a multimodal hub (*Bimodal Hubs < Shallow Multimodal Hub*; t(2087.349)=-9.550, p<.001) and shortcut connections (*Deep Multimodal Hub< Multimodal Hub-plus-Shortcut*; t(430.907)=-2.803, p<.05; *Convergent Hubs* *< Convergent Hubs-plus-Shortcut*; t(458.843)=-3.882, p<.001). Depth had no significant effect (*Shallow Multimodal Hub vs. Deep Multimodal Hub*; t(958)=-1.534, p=.125), whilst hierarchical convergence significantly decreased similarity to the unimodal structure (*Deep Multimodal Hub vs. Convergent Hubs*; t(382.124)=8.523, p<.001; *Multimodal Hub-plus-Shortcut vs. Convergent Hubs-plus-Shortcut*; t(414.305)=4.393, p<.001). Thus, in simulations without control the building blocks identified to promote conceptual abstraction appear to be true for all these structures, and may not critically depend on the integration across modalities.

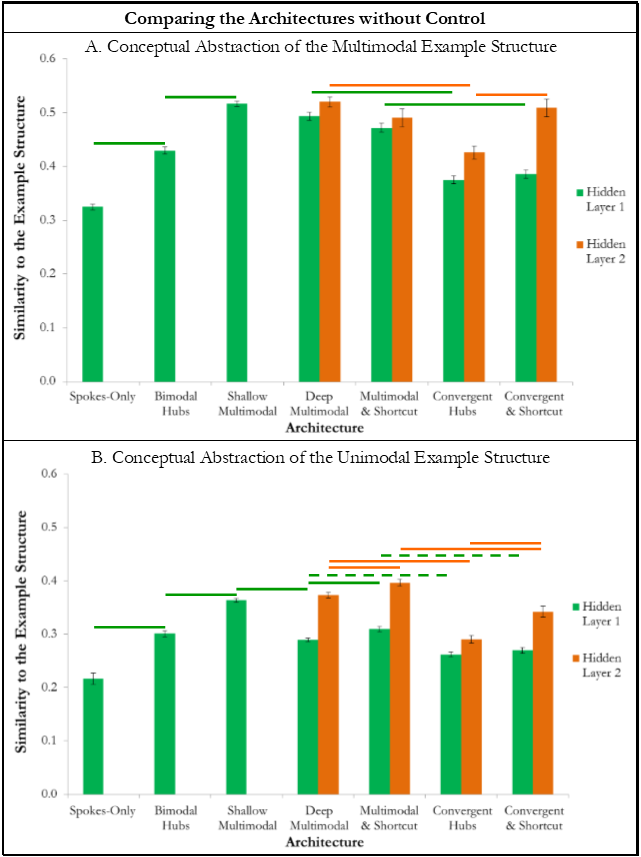

*Supplementary Figure 5. Comparing how well the architectures without control learnt the orthogonal multimodal and unimodal example structures. A. The similarity between the multimodal example structure and the representations in Hidden Layer 1 (green) and Hidden Layer 2 (orange) of each architecture. B. The similarity between the unimodal example structure and the representations in Hidden Layer 1 (green) and Hidden Layer 2 (orange) of each architecture. A&B. The bars show the mean similarity value across the different runs of the model (error bars show standard error). Significant differences in the planned contrasts are highlighted with a line (p<.05). Significant contrasts in Hidden Layer 1 are shown in green, whereas Hidden Layer 2 differences are shown in orange. Where Hidden Layer 1 and 2 are contrasted between the shallow and deep multimodal hub models, a black line reflects significance. If a contrast is significant, yet the size-matched equivalent is not, the line is dashed to highlight that this difference may be induced through the comparison of different numbers of units.*

The same finding is apparent in the simulations with control, where extraction of the context-independent multimodal structure differs based on architecture (Supplementary Figure 6.A; Hidden Layer 1; F(6,2393)=278.121, p<.001; Hidden Layer 2; F(3,316)=266.094, p<.001; all significant contrasts have p<.001) and are improved by the inclusion of a hub (*Spokes-Only < Bimodal Hubs*; t(639.345)=-24.104) the hub being multimodal (*Bimodal Hubs < Shallow Multimodal Hub*; t(718)=-34.873), depth (*Shallow Multimodal Hub < Deep Multimodal Hub*; t(103.307)=-10.067) and shortcut connections (*Deep Multimodal Hub <Multimodal Hub-plus-Shortcut*; t(140.239)=-7.007, p<.001; *Convergent Hubs* *< Convergent Hubs-plus-Shortcut*; t(83.761)=-16.929, p<.001) yet reduced by hierarchical convergence (*Deep Multimodal Hub > Convergent Hubs*; t(103.150)=39.484, p<.001; *Multimodal Hub-plus-Shortcut > Convergent Hubs-plus-Shortcut*; t(134.009)=5.795, p<.001).

The similarity with the context-independent unimodal example structures also varied by architecture (Supplementary Figure 6.B.; Hidden Layer 1; F(6,7193)=439.144, p<.001; Hidden Layer 2; F(3,956)=416.942, p<.001; all significant contrasts have p<.001) and depended on the same building blocks (*Spokes-Only < Bimodal Hubs*; t(2158)=-14.247, p<.001; *Bimodal Hubs < Shallow Multimodal Hub;* t(1144.701)=-44.763, p<.001; *Shallow Multimodal Hub < Deep Multimodal Hub*; t(324.658)=-14.840, p<.001; *Convergent Hubs* *< Convergent Hubs-plus-Shortcut*; t(252.840)=-19.182, p<.001; *Deep Multimodal Hub > Convergent Hubs*; t(103.150)=39.484, p<.001; *Multimodal Hub-plus-Shortcut > Convergent Hubs-plus-Shortcut*; t(134.009)=5.795, p<.001). Thus, both the multimodal and unimodal structures showed precisely the same pattern as the full structure; a controlled semantic system should employ a single, deep multimodal hub with shortcut connections in order to distil context-independent representations.

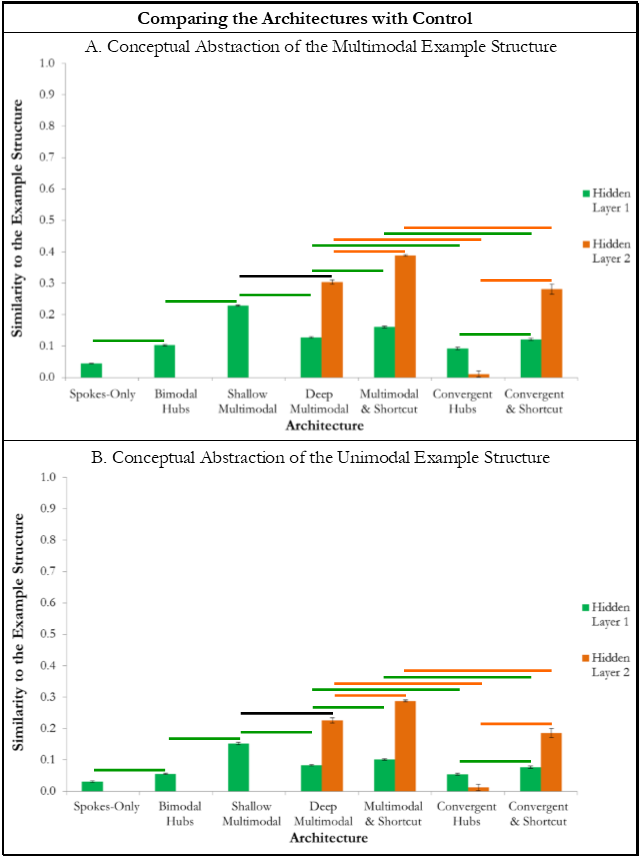

*Supplementary Figure 6. Comparing how well the architectures with control learnt the orthogonal multimodal and unimodal example structures. A. The similarity between the multimodal example structure and the representations in Hidden Layer 1 (green) and Hidden Layer 2 (orange) of each architecture. B. The similarity between the unimodal example structure and the representations in Hidden Layer 1 (green) and Hidden Layer 2 (orange) of each architecture. A&B. The bars show the mean similarity value across the different runs of the model (error bars show standard error). Significant differences in the planned contrasts are highlighted with a line (p<.05). Significant contrasts in Hidden Layer 1 are shown in green, whereas Hidden Layer 2 differences are shown in orange. Where Hidden Layer 1 and 2 are contrasted between the shallow and deep multimodal hub models, a black line reflects significance. If a contrast is significant, yet the size-matched equivalent is not, the line is dashed to highlight that this difference may be induced through the comparison of different numbers of units.*

Supplementary Note 8: Varying the Sparsity of Shortcut Connections

As the inclusion of shortcut connections was found to be beneficial, the effect of varying the amount of shortcut connections in the *Multimodal Hub-plus-Shortcut* architecture was assessed. The proportion of connections included varied from no shortcut connections (i.e., the *Deep Multimodal Hub* architecture), to 1 in 3 possible shortcut connections (432 connections). Intermediate amounts of shortcut connections included 1 in 36 (36 connections), 1 in 24 (i.e., the original *Multimodal Hub-plus-Shortcut* architecture with 54 connections), 1 in 16 (81 connections), 1 in 12 (108 connections) and 1 in 6 (216 connections). All models were matched for number of total connections; therefore those with more shortcut connections had fewer connections from Hidden Layer 1 to Hidden Layer 2. This meant varying the proportion of connections between these layers from approximately 0.671 to approximately 0.385. All other details were identical to the main simulations. Contrasts were performed comparing each consecutive pair of models (to assess the effect of increasing the number of connections in that step) and comparing the version with no shortcut connections with the version with 1 in 3 (to assess the significance of the most extreme increase in shortcut connections tested, in case the difference at each consecutive step was too subtle for statistical significance). The results were Bonferroni corrected for 7 contrasts. Supplementary Figure 7.A. shows the gains in similarity on the conceptual abstraction score at each step, Supplementary Figure 7.B. the similarity to the Context-Only structure, and Supplementary Figure 7.C. the change in number of epochs taken to reach the learning criterion.

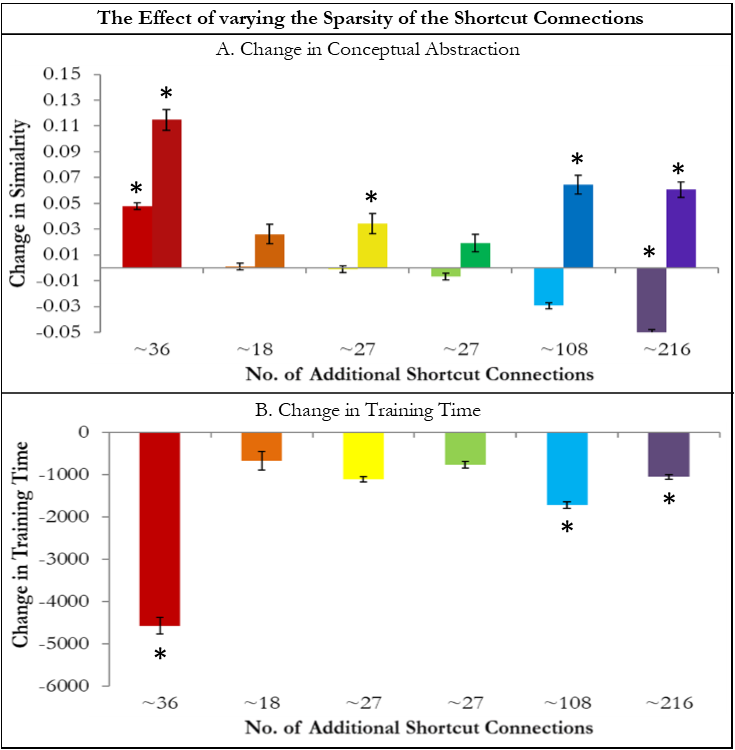

*Supplementary Figure 7. The effect of varying the number of shortcut connections. The gain in each measure is plotted against the number of additional shortcut connections in each model version. Each column is contrasted with the prior one, with the first being compared to the Deep Multimodal architecture as this has no shortcut connections. A. The change in similarity to the Context-Independent example structure in Hidden Layer 1 (lighter bars) and Hidden Layer 2 (darker bars). Significant changes (p<.05) are highlighted with an asterisk. B. The change in time taken (epochs) to learn the similarity structure. Significant changes (p<.05) are highlighted with an asterisk.*

Each successive step led to a numeric increase in the conceptual abstraction score although not all reached significance (F(6, 553)=181.173, p<.001; *Deep Multimodal Hub < 1 in 36;* t(158)=-9.099, p<.001; *1 in 36 ~= 1 in 24;* t(158)=-2.362, p=.136; *1 in 24 < 1 in 16;* t(158)=-3.194, p<.05; *1 in 16~= 1 in 12;* t(158)=-1.882, p=.432; *1 in 12 < 1 in 6;* t(158)=-6.388, p<.001; *1 in 6 < 1 in 3;* t(150.266)=-6.437, p<.001; *Deep Multimodal Hub < 1 in 3; t*(130.532)=-28.355, p<.001*).* Similarity to the context-independent semantic representation structure in Hidden Layer 1 was initially increased and then decreased (F(6, 1673)=206.045, p<.001; *Deep Multimodal Hub < 1 in 36;* t(450.339)=-15.078, p<.001; *1 in 36 ~= 1 in 24;* t(478)=-0.318, p=1; *1 in 24 ~= 1 in 16;* t(478)=0.277, p=1; *1 in 16 ~= 1 in 12;* t(478)=1.909, p=.400; *1 in 12 > 1 in 6;* t(478)=8.769, p<.001; *1 in 6 > 1 in 3;* t(460.768)=16.593, p<.001; *Deep Multimodal Hub > 1 in 3;* t(478)=13.992, p<.001*).* The time taken to learn the concepts is decreased when the number of shortcut connections is increased (F(6,98)=98.072, p<.001; *Deep Multimodal Hub > 1 in 36;* t(28)=7.619, p<.001; *1 in 36 ~= 1 in 24;* t(28)=1.059, p=1; *1 in 24 ~= 1 in 16;* t(28)=2.078, p=.329; *1 in 16 ~= 1 in 12;* t(28)=2.252, p=.227; *1 in 12 > 1 in 6;* t(28)=5.683, p<.001; *1 in 6 > 1 in 3;* t(23.201)=4.628, p<.001; *Deep Multimodal Hub > 1 in 3;* t(16.191)=22.473, p<.001).

Thus, the core metrics suggest that it is optimal to increase the number of shortcut connections (at least up to 1 in 3 connections). However, the greatest positive effect on these metrics occurs with the initial addition of merely 36 connections (i.e., between the *Deep Multimodal Hub* architecture and *1 in 36* connections). This addition of a small number of connections results in the greatest gain and each subsequent increase leads to diminishing returns. Thus, as the brain must balance improved performance and quicker learning times with the harsh packing and metabolic constraints, it may be that a relatively small proportion of connections inducing a large change is the optimal solution. As the ideal balance between the cognitive improvement and minimising the packing and metabolic constraints cannot be determined here, all analyses continue to use the original sparsity level.

Supplementary Note 9: Similarity to the Context-Only and Context-Sensitive Structures

The different context-sensitive architectures may also be assessed on how strongly the regions represent the Context-Only structure, although this does not demonstrate the aptitude of a given model. This is the corollary of the context-independent semantic representation structure and is therefore, often inversely-related although this is not always the case. Similarity to the Context-Only structure varied by architecture in both Hidden Layer 1 (F(6, 2393)=215.479, p<.001) and Hidden Layer 2 (F(3, 316)=462.595, p<.001), shown in Supplementary Figure 8.A. Similarity to the Context-Only structure was significantly reduced by the presence of a multimodal, but not a bimodal, hub (*Spokes-Only ~= Bimodal Hubs*; t(667.833)=2.165, p=.215; *Bimodal Hubs > Shallow Multimodal Hub*; t(696.732)=23.225, p<.001). Depth had no effect (*Shallow Multimodal Hub ~=Deep Multimodal Hub*; t(86.396)=-0.871, p=.386) whereas shortcut connections led to a significant reduction in the representation of the Context-Only structure (*Deep Multimodal Hub > Multimodal Hub-plus-Shortcut*; t(158)=9.589, p<.001; *Convergent Hubs* *> Convergent Hubs-plus-Shortcut*; t(122.016)=27.593, p<.001). Hierarchical convergence increased the similarity with the Context-Only structure (*Deep Multimodal Hub<* *Convergent Hubs*; t(131.743)=-27.937, p<.001; *Multimodal Hub-plus-Shortcut* *< Convergent Hubs-plus-Shortcut*; t(144.271)=-6.815, p<.001).

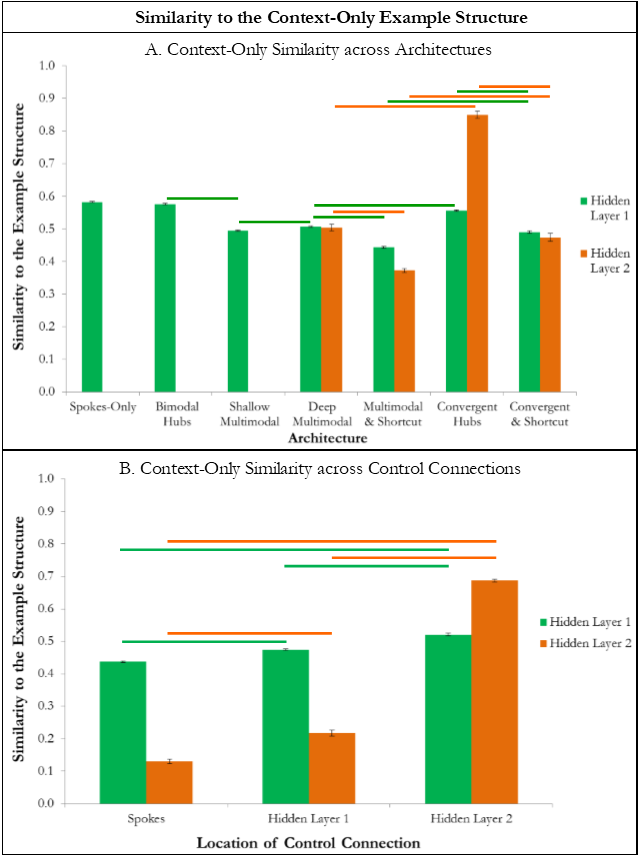

*Supplementary Figure 8. Similarity of the models representations to the Context-Only example structure. A. Similarity across the architectures in in Hidden Layer 1 (green) and Hidden Layer 2 (orange) of each architecture. B. Similarity to the Context-Only example structure in Hidden Layer 1 (green) and Hidden Layer 2 (orange) with the control signal connected to the Spokes Layer, Hidden Layer 1 or Hidden Layer 2. A&B. The bar shows the mean similarity value across the different runs of the model (bars show standard error). Significant differences in the planned contrasts are highlighted with a line (p<.05). Significant contrasts in Hidden Layer 1 are shown in green, whereas Hidden Layer 2 differences are shown in orange.*

Similarity to the Context-Only examples structure was also assessed for the simulations where the control signal was connected to the Spoke Layer, Hidden Layer 1 or Hidden Layer 2 (see Supplementary Figure 8.B.). Whilst similarity to the Context-Only structure in Hidden Layer 2 is typically low, it increases dramatically if the control signal is directly connected to this region (Spokes Layer < Hidden Layer 2; t(158)=-57.187, p<.001; Hidden Layer 1 *<* Hidden Layer 2; t(148.199)=-39.662, p<.001). This is accompanied by an increase in similarity in Hidden Layer 1 (Spokes Layer *<* Hidden Layer 2; t(453.499)=-25.259, p<.001; Hidden Layer 1 *<* Hidden Layer 2; t(478)=-15.832, p<.001). A smaller, yet significant, effect was identified between the Spokes Layer and Hidden Layer 1 connections; connecting the control units to Hidden Layer 1 led to significantly higher similarity with the Context-Only structure in both Hidden Layer 1 and 2 (Hidden Layer 1; Spokes Layer *<* Hidden Layer 1; t(457.575)=-11.134, p<.001; Hidden Layer 2; Spokes Layer *<* Hidden Layer 1; t(158)=-7.600, p<.001).

In combination the Context-Only and Context-Independent example structures, form the full structure engaged in the task, the Context-Sensitive example structure. This structure reflects the similarity between concepts in a given task context, which all the models must learn in order to perform the task and is therefore not our core focus. However, some architectures learn representations that reflect this structure more or less closely (Supplementary Figure 9; Hidden Layer 1; F(6,2393)=456.802, p<.001; Hidden Layer 2; F(3,316)=86.530, p<.001). Similarity to the Context-Sensitive structure is increased with the presence of a hub (*Spokes-Only < Bimodal Hubs*; t(610.452)=-18.755, p<.001) and, in particular, a multimodal hub (*Bimodal Hubs < Shallow Multimodal Hub*; t(618.788)=-35.948, p<.001). Depth increases this similarity (*Shallow Multimodal Hub* < *Deep Multimodal Hub*; t(100.162)=-18.766, p<.001), whilst it is reduced by hierarchical convergence (*Deep Multimodal Hub > Convergent Hubs*; t(148.075)=15.435, p<.001; *Multimodal Hub-plus-Shortcut > Convergent Hubs-plus-Shortcut*; t(147.894)=8.359, p<.001). Shortcut connections reduced this similarity only when present without hierarchical convergence (*Deep Multimodal Hub > Multimodal Hub-plus-Shortcut*; t(158)=3.165, p<.01; *Convergent Hubs ~= Convergent Hubs-plus-Shortcut*; t(123.451)=-0.187, p=1).

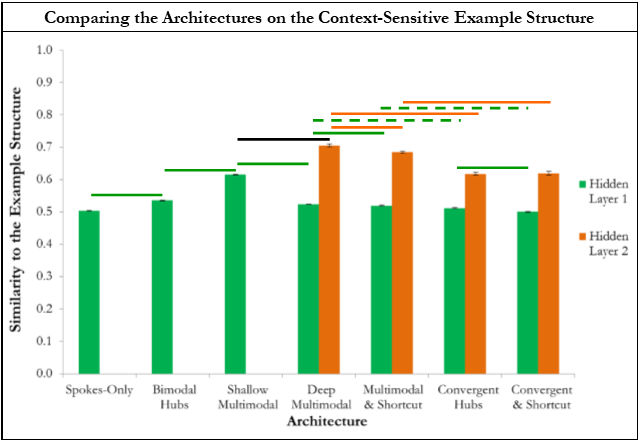

*Supplementary Figure 9. Comparing how well the architectures with control learnt the Context-Sensitive example structure in Hidden Layer 1 (green) and Hidden Layer 2 (orange) of each architecture. The bars show the mean similarity value across the different runs of the model (error bars show standard error). Significant differences in the planned contrasts are highlighted with a line (p<.05). Significant contrasts in Hidden Layer 1 are shown in green, whereas Hidden Layer 2 differences are shown in orange. Where Hidden Layer 1 and 2 are contrasted between the shallow and deep multimodal hub models, a black line reflects significance. If a contrast is significant, yet the size-matched equivalent is not, the line is dashed to highlight that this difference may be induced through the comparison of different numbers of units.*

Supplementary Methods 1: Matching the Number of Connections across Architectures

To match the number of connections between architectures all (except the *Bimodal Hubs* architecture which had the fewest possible connections) had some sparse connections. These were connections within Hidden Layer 1 for the *Spokes-Only,* *Shallow Multimodal Hub*, *Deep Multimodal Hub* and *Multimodal Hub-plus-Shortcut* architectures (with each connection applied at a probability of 0.28, 0.333333, 0.5 and 0.5 respectively) and between the hidden layers for the *Deep Multimodal Hub,* *Multimodal Hub-plus-Shortcut,* *Convergent Hubs* and *Convergent Hubs-plus-Shortcut* architectures (with each connection applied at a probability of 0.670634920634921, 0.634920653, 0.476190476 and 0.44047619047619).

Supplementary Methods 2: Simulation Code for LENS

Simulations without Control (uncomment where suggested in red for each architecture and to add or remove the requirement for control)

addNet hier -i 6 -t 4 CONTINUOUS

addGroup m1 12 INPUT

addGroup m2 12 INPUT

addGroup m3 12 INPUT

#TURN ON FOR SIMULATIONS WITH CONTROL

#addGroup Control 3 INPUT

addGroup m1rep 12 OUTPUT CROSS_ENTROPY -BIASED

addGroup m2rep 12 OUTPUT CROSS_ENTROPY -BIASED

addGroup m3rep 12 OUTPUT CROSS_ENTROPY -BIASED

#TURN ON FOR SHALLOW ARCHITECTURES

#addGroup m1m2 10

#addGroup m2m1 10

#addGroup m1m3 10

#addGroup m3m1 10

#addGroup m2m3 10

#addGroup m3m2 10

#TURN ON FOR DEEP ARCHITECTURES

#addGroup m1m2 7

#addGroup m2m1 7

#addGroup m1m3 7

#addGroup m3m1 7

#addGroup m2m3 7

#addGroup m3m2 7

#addGroup atl 18

#Set biases on feature units to -3

connectGroups bias m1rep -m -3 -r 0

connectGroups bias m2rep -m -3 -r 0

connectGroups bias m3rep -m -3 -r 0

#Make sure biases on feature units can't learn

freezeWeights -g {m1rep m2rep m3rep} -t bias

#Connect input features to corresponding feature units with

#a weight of +6, so net input for active features will be

#+3 and for inactive features will be -3

connectGroups m1 m1rep -p ONE_TO_ONE -m 6 -r 0

connectGroups m2 m2rep -p ONE_TO_ONE -m 6 -r 0

connectGroups m3 m3rep -p ONE_TO_ONE -m 6 -r 0

#Self-connections among feature units

connectGroups m1rep m1rep

connectGroups m2rep m2rep

connectGroups m3rep m3rep

#TURN ON FOR SIMULATIONS WITH CONTROL

#connectGroups Control m1rep

#connectGroups Control m2rep

#connectGroups Control m3rep

#connectGroups Control m1m2

#connectGroups Control m1m3

#connectGroups Control m2m1

#connectGroups Control m2m3

#connectGroups Control m3m1

#connectGroups Control m3m2

#Self-connections among hidden unit layers including connections within region, creating hubs #and resource matching

#TURN ON FOR SPOKES-ONLY ARCHITECTURE ONLY

#connectGroups m1m2 m1m2 -p RANDOM -s 0.28

#connectGroups m1m3 m1m3 -p RANDOM -s 0.28

#connectGroups m2m1 m2m1 -p RANDOM -s 0.28

#connectGroups m2m3 m2m3 -p RANDOM -s 0.28

#connectGroups m3m1 m3m1 -p RANDOM -s 0.28

#connectGroups m3m2 m3m2 -p RANDOM -s 0.28

#connectGroups m1m2 m1m3 -bi -p RANDOM -s 0.28

#connectGroups m2m1 m2m3 -bi -p RANDOM -s 0.28

#connectGroups m3m1 m3m2 -bi -p RANDOM -s 0.28

#TURN ON FOR BIMODAL HUBS AND CONVERGENT HUBS AND CONVERGENT HUB-PLUS-SHORTCUT ARCHITECTURES ONLY

#connectGroups m1m2 m1m2

#connectGroups m1m3 m1m3

#connectGroups m2m1 m2m1

#connectGroups m2m3 m2m3

#connectGroups m3m1 m3m1

#connectGroups m3m2 m3m2

#connectGroups m1m2 m2m1 -bi

#connectGroups m1m3 m3m1 -bi

#connectGroups m2m3 m3m2 -bi

#TURN ON FOR SHALLOW MULTIMODAL HUB ARCHITECTURE ONLY

#connectGroups m1m2 m1m2 -p RANDOM -s 0.333333

#connectGroups m1m3 m1m3 -p RANDOM -s 0.333333

#connectGroups m2m1 m2m1 -p RANDOM -s 0.333333

#connectGroups m2m3 m2m3 -p RANDOM -s 0.333333

#connectGroups m3m1 m3m1 -p RANDOM -s 0.333333

#connectGroups m3m2 m3m2 -p RANDOM -s 0.333333

#connectGroups m1m2 m2m1 -bi -p RANDOM -s 0.333333

#connectGroups m1m2 m1m3 -bi -p RANDOM -s 0.333333

#connectGroups m1m2 m2m3 -bi -p RANDOM -s 0.333333

#connectGroups m1m2 m3m2 -bi -p RANDOM -s 0.333333

#connectGroups m1m2 m3m1 -bi -p RANDOM -s 0.333333

#connectGroups m2m1 m1m3 -bi -p RANDOM -s 0.333333

#connectGroups m2m1 m2m3 -bi -p RANDOM -s 0.333333

#connectGroups m2m1 m3m2 -bi -p RANDOM -s 0.333333

#connectGroups m2m1 m3m1 -bi -p RANDOM -s 0.333333

#connectGroups m1m3 m3m1 -bi -p RANDOM -s 0.333333

#connectGroups m1m3 m3m2 -bi -p RANDOM -s 0.333333

#connectGroups m1m3 m2m3 -bi -p RANDOM -s 0.333333

#connectGroups m3m1 m3m2 -bi -p RANDOM -s 0.333333

#connectGroups m3m1 m2m3 -bi -p RANDOM -s 0.333333

#connectGroups m3m2 m2m3 -bi -p RANDOM -s 0.333333

#TURN ON FOR DEEP ARCHITECTURES ONLY

#connectGroups atl atl

#TURN ON FOR DEEP MULTIMODAL HUB & MULTIMODAL HUB-PLUS-SHORTCUT ARCHITECTURES ONLY

#connectGroups m1m2 m1m2 -p RANDOM -s 0.5

#connectGroups m1m3 m1m3 -p RANDOM -s 0.5

#connectGroups m2m1 m2m1 -p RANDOM -s 0.5

#connectGroups m2m3 m2m3 -p RANDOM -s 0.5

#connectGroups m3m1 m3m1 -p RANDOM -s 0.5

#connectGroups m3m2 m3m2 -p RANDOM -s 0.5

#connectGroups m1m2 m1m3 -bi -p RANDOM -s 0.5

#connectGroups m2m1 m2m3 -bi -p RANDOM -s 0.5

#connectGroups m3m1 m3m2 -bi -p RANDOM -s 0.5

#TURN ON FOR DEEP MULTIMODAL HUB ARCHITECTURE ONLY

#connectGroups m1m2 atl -bi -p RANDOM -s 0.670634920634921

#connectGroups m2m1 atl -bi -p RANDOM -s 0.670634920634921

#connectGroups m1m3 atl -bi -p RANDOM -s 0.670634920634921

#connectGroups m3m1 atl -bi -p RANDOM -s 0.670634920634921

#connectGroups m2m3 atl -bi -p RANDOM -s 0.670634920634921

#connectGroups m3m2 atl -bi -p RANDOM -s 0.670634920634921

#TURN ON FOR MULTIMODAL HUB-PLUS-SHORTCUT ARCHITECTURE ONLY

#connectGroups m1m2 atl -bi -p RANDOM -s 0.634920635

#connectGroups m2m1 atl -bi -p RANDOM -s 0.634920635

#connectGroups m1m3 atl -bi -p RANDOM -s 0.634920635

#connectGroups m3m1 atl -bi -p RANDOM -s 0.634920635

#connectGroups m2m3 atl -bi -p RANDOM -s 0.634920635

#connectGroups m3m2 atl -bi -p RANDOM -s 0.634920635

#TURN ON FOR CONVERGENT HUB ARCHITECTURE ONLY

#connectGroups m1m2 atl -bi -p RANDOM -s 0.476190476

#connectGroups m2m1 atl -bi -p RANDOM -s 0.476190476

#connectGroups m1m3 atl -bi -p RANDOM -s 0.476190476

#connectGroups m3m1 atl -bi -p RANDOM -s 0.476190476

#connectGroups m2m3 atl -bi -p RANDOM -s 0.476190476

#connectGroups m3m2 atl -bi -p RANDOM -s 0.476190476

#TURN ON FOR CONVERGENT HUB-PLUS-SHORTCUT ARCHITECTURE ONLY

#connectGroups m1m2 atl -bi -p RANDOM -s 0.44047619047619

#connectGroups m2m1 atl -bi -p RANDOM -s 0.44047619047619

#connectGroups m1m3 atl -bi -p RANDOM -s 0.44047619047619

#connectGroups m3m1 atl -bi -p RANDOM -s 0.44047619047619

#connectGroups m2m3 atl -bi -p RANDOM -s 0.44047619047619

#connectGroups m3m2 atl -bi -p RANDOM -s 0.44047619047619

#shortcut connections

#TURN ON FOR MULTIMODAL HUB-PLUS-SHORTCUT AND CONVERGENT HUB-PLUS-SHORTCUT ARCHITECTURES ONLY

#connectGroups m1rep atl -bi -p RANDOM -s 0.041666667

#connectGroups m2rep atl -bi -p RANDOM -s 0.041666667

#connectGroups m3rep atl -bi -p RANDOM -s 0.041666667

#TURN ON FOR BIMODAL HUBS & CONVERGENT HUBS & CONVERGENT HUBS-PLUS-SHORTCUT ARCHITECTURES ONLY

#Connections among first layer hidden units (separate paths)

#connectGroups m1m2 m2m1 -bi

#connectGroups m1m3 m3m1 -bi

#connectGroups m2m3 m3m2 -bi

#TURN ON FOR SPOKES-ONLY ARCHITECTURE ONLY

#direct paths amongst surface representations:

#connectGroups m1rep m2rep -bi

#connectGroups m1rep m3rep -bi

#connectGroups m2rep m3rep –bi

#connect feature units to their paired hidden layers

connectGroups m1rep m1m2 -bi

connectGroups m1rep m1m3 -bi

connectGroups m2rep m2m1 -bi

connectGroups m2rep m2m3 -bi

connectGroups m3rep m3m1 -bi

connectGroups m3rep m3m2 –bi

#TURN ON FOR SIMULATIONS WITHOUT CONTROL

loadExamples FIXED_EXAMPLES_USING_hierarchical_model.ex -exmode PER -set train

loadExamples FIXED_EXAMPLES_USING_hierarchical_model.ex -exmod ORD -set test

#TURN ON FOR SIMULATIONS WITH CONTROL

#loadExamples hier_examples_basic_control.ex -exmode PER -set train

#loadExamples hier_examples_basic_control.ex -exmod ORD -set test

setObj learningRate 0.001

setObj numUpdates 500000

setObj weightDecay 0.0001

#Training will stop when output units are less than 0.2 from target:

setObj trainGroupCrit 0.2

train -a steepest
